# Structure of *Bacillus cereus* ZmaM, a peptidase-ABC transporter that secretes and activates zwittermicin A

**DOI:** 10.64898/2026.10.31.763113

**Authors:** Gerardo E. Zavala, Jose A. Velilla, Richard M. Walsh, Mit Patel, Karissa Mennes, Wasan Rafat, Miguel A. Aguilar Ramos, Martin Daniel-Ivad, Fernando H. Ramírez-Guadiana, David Z. Rudner, Emily P. Balskus, Rachelle Gaudet

## Abstract

Bacteria engage in microbial warfare by secreting toxic molecules. These toxins can serve as therapeutics or drug discovery starting points, motivating investigations of their production. Bacteria that produce these toxins must protect themselves, and a common strategy is to synthesize an inactive prodrug that is activated upon secretion. Some prodrugs are capped with *N*-acyl-ᴅ-asparagine motifs, which are removed by two types of prodrug peptidases. Type I peptidases are membrane-bound S12 hydrolases; type II peptidases have an additional C-terminal half-ABC transporter module. Recent structures of a type I peptidase revealed how its transmembrane domain completes the substrate-binding site, but little is known about type II peptidases. Here, we show that the *B. cereus* type II peptidase ZmaM is responsible for both the export of prezwittermicin and its activation into zwittermicin. The structure of homodimeric ZmaM reveals a central type IV ABC transporter flanked by peptidase modules. The periplasmic hydrolase domains interact over the transporter’s open outer vestibule. Mutations to intermodule interfaces impact both peptidase and ATP hydrolysis activities, indicating the peptidase and transporter modules are interdependent. Together, our findings reveal an evolutionary strategy to coordinate toxin activation with its export and inform future efforts to harness these systems in biotechnology.

## Introduction

Bacteria compete for scarce environmental resources through microbial warfare. In one approach, they produce and secrete toxins with antifungal, antiprotist, and/or antibacterial activities. These specialized metabolites are of interest for their potential use as medicines and pesticides^1^. Because these toxins typically target conserved and essential machinery in competing microbes, they pose a risk to the producing bacteria, which have evolved strategies to mitigate this risk. A classic example is degradation of β-lactam antibiotics mediated by β-lactamases^2^. Another strategy is to synthesize a nontoxic precursor and activate it once safe to do so, as in the case of toxins capped by an *N*-acyl-ᴅ-asparagine prodrug motif. These prodrug toxins have garnered interest because they encompass medically and ecologically relevant molecules^3,4^, including colibactin^5^, xenocoumacin^3^, paenilamicin^6^, amicoumacins^7^, edeines^8^, and the broad-spectrum antibiotic, antifungal, and antiprotist zwittermicin A^9,10^. Zwittermicin has been found in the rhizosphere of alfalfa and the skin microbiota of amphibians, protecting both organisms from fungal pathogens among other microbes^11,12^. Its broad activity spectrum and prevalence in nature has prompted the use of zwittermicin-producing microbes as biocontrol agents and pesticides^13^.

*N*-acyl-ᴅ-asparagine prodrug toxins are produced by hybrid non-ribosomal peptide synthase and polyketide synthase biosynthetic gene clusters (BGCs)^3^. The *N*-acyl-ᴅ-asparagine motif is generated and installed early in the cytosolic synthesis pathway, blocking functional groups essential for toxin activity^14^. The completed prodrug is exported to the periplasm where the prodrug motif is removed enzymatically^4,14^. This removal is accomplished by membrane-bound S12 hydrolases known as prodrug peptidases^3,4^. These enzymes form two groups: type I peptidases comprise an N-terminal periplasmic hydrolase domain and a C-terminal three-helix transmembrane domain (TMD); type II peptidases have a type I-like peptidase module followed by a C-terminal ATP-binding-cassette (ABC) half-transporter module. Importantly, BGCs with type II prodrug peptidases do not encode another transporter, suggesting that type II peptidases both export and activate the nontoxic prodrug^14^. Understanding the mechanism of these final steps in toxin synthesis is important because the periplasmic peptidase is an attractive target for modulating toxin production^15^, and both transport and activation could be manipulated in bioengineering efforts to leverage prodrug toxin BCGs^4^.

The best characterized type I prodrug peptidase is ClbP, the peptidase responsible for activating colibactin, a genotoxin associated with colorectal cancer^16–18^. We recently reported structures of full-length ClbP, revealing how its TMD completes the substrate-binding site^17,18^, which explains the essential role of the TMD in colibactin production^19^. ClbP is an obligate homodimer, with an extensive interface formed by the two hydrolase domains but no intersubunit interactions mediated by the two TMDs. The two active sites of the ClbP dimer can accommodate the pseudodimeric precolibactin, suggesting that the dimer could help insure near-simultaneous activation of colibactin’s dual warheads^18^.

Despite their unusual architecture, type II prodrug peptidases are structurally and biochemically underexplored compared to their type I counterparts. Moreover, sequence mining for BGCs containing prodrug peptidases identified many that produce unknown metabolites^4^. These BGCs are in marine and soil bacteria which have long been sources of new antibiotics, and in species populating microbiomes where they may play roles in host-microbe interactions^4^. Understanding how prodrug peptidases function has the potential to inform discovery of inhibitors that selectively suppress the production of harmful toxins or bioengineering of specialized metabolite production, export, and activation.

We chose ZmaM, a component of the zwittermicin BCG found in *Bacillus cereus* and *Bacillus thuringiensis* strains, as a model type II prodrug peptidase. ZmaM has been hypothesized to export and activate the precursor prezwittermicin^14,20^. The complete zwittermicin BCG and culture conditions that promote zwittermicin production in the readily culturable *B. cereus* UW85 strain have been identified^21–23^. Here, we genetically engineer *B. cereus* strains and demonstrate that ZmaM both exports and activates prezwittermicin. Purified, detergent-solubilized ZmaM produces zwittermicin from prezwittermicin isolated from a *B. cereus* strain with a *zmaM* deletion. Our cryogenic electron microscopy (cryo-EM) structure of the ZmaM homodimer shows its ABC transporter modules in an outward-open conformation. The two peptidase modules—dimerized in a manner unexpectedly analogous to the ClbP homodimer—straddle the transporter such that its substrate-exit vestibule faces the two peptidase active sites. Structure-guided point mutations to various interdomain interfaces impaired both peptidase and ATPase activity. Overall, our biochemical and structural data indicate that the peptidase and transporter modules of ZmaM are interdependent and structurally organized such that their activities are coordinated.

## Results

### ZmaM secretes and activates zwittermicin

ZmaM is the only transporter and prodrug peptidase encoded within the zwittermicin BGC (Fig. 1a). Because the BGC is sufficient for zwittermicin production^20,24^, ZmaM has been hypothesized to both transport and activate zwittermicin (Fig. 1b)^14,20^. However, this hypothesis has not been experimentally tested. To investigate ZmaM’s *in vivo* activity, we generated a ZmaM-deficient strain of *B. cereus* UW85 (Δ*zmaM)* by allelic exchange (Supplementary Fig. 1)^25^. This deletion retained 3 N-terminal and 7 C-terminal codons to preserve the reading frame and minimize polar effects. We compared the accumulation of prezwittermicin and zwittermicin in spent media or methanol-extracted UW85 wildtype (WT) and Δ*zmaM* cell pellets by liquid chromatography–mass spectrometry (LC-MS). Δ*zmaM*, but not WT, accumulated prezwittermicin both intracellularly and extracellularly, and extracellular zwittermicin accumulation was reduced but not completely eliminated in Δ*zmaM* (Supplementary Fig. 2).

**Figure 1.**
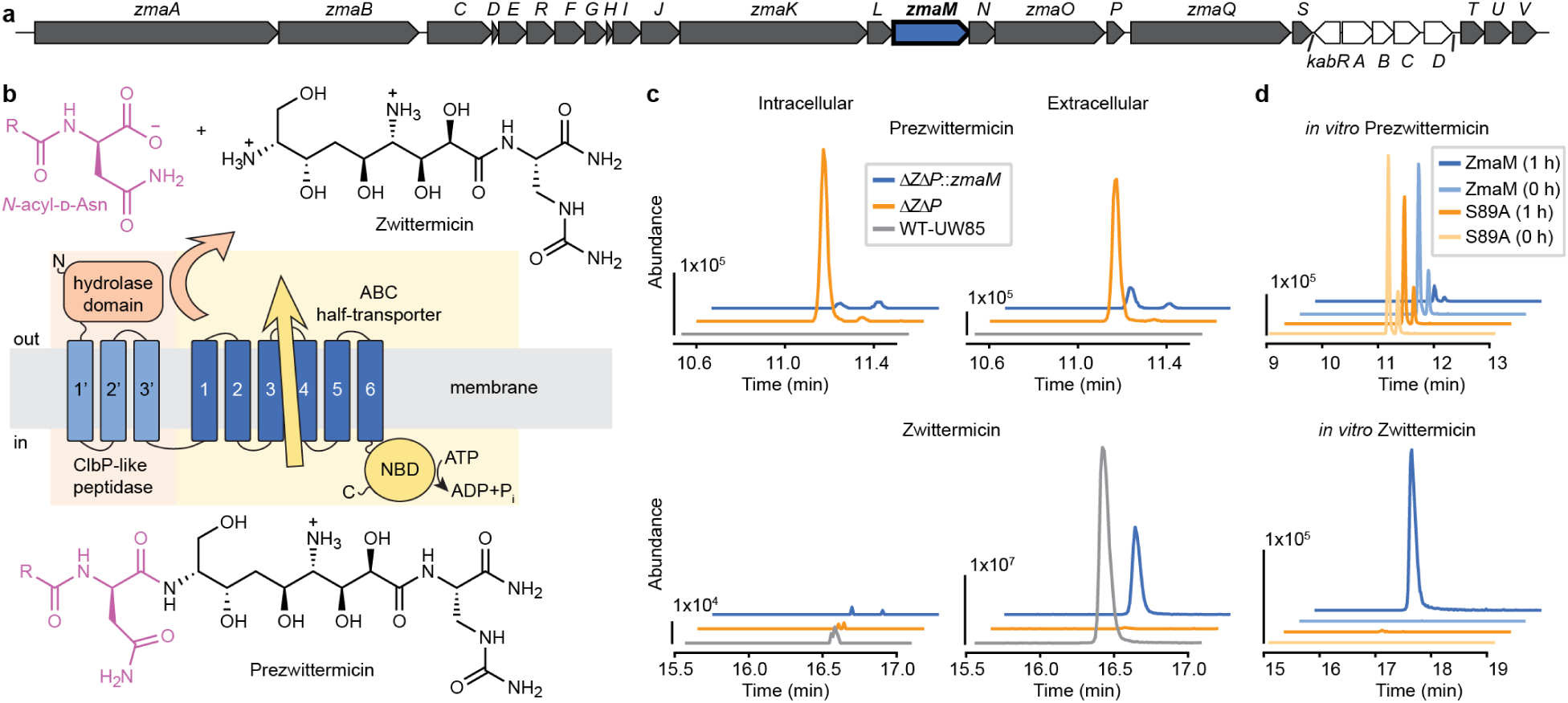
ZmaM activates and secretes zwittermicin in *B. cereus* UW85. **a**, The zwittermicin-producing BGC from *B. cereus* UW85 (∼63 kilobases), with the *zmaM* gene in blue. The unrelated kanosamine BGC is in white. **b,** Reaction scheme for prezwittermicin export and activation by ZmaM. Prodrug moiety is pink. The domain organization and membrane topology of ZmaM is illustrated in the center. The prezwittermicin and zwittermicin protonation states are shown as predicted for pH 7 using MolGpka^65^. **c,** LC-MS Extracted Ion Chromatograms (EICs) showing the abundance of prezwittermicin (top, EIC for [M+H]^+^ = 693.4141) and zwittermicin (bottom, EIC for [M+H]^+^ = 315.2278) in organic extracts of cell pellets (intracellular; left) and spent media (extracellular; right). Y-axis scales differ between plots but are consistent within each plot. **d,** EICs for *in vitro* activation of prezwittermicin by purified samples of ZmaM and the peptidase-deficient S89A variant. The data in panels (c) and (d) are representative traces of n = 2 biological replicates.

We hypothesized that this residual zwittermicin activation by Δ*zmaM* resulted from an alternative pathway. We identified a type I prodrug peptidase gene in the UW85 genome (name *proP*), which is not part of any identifiable BGC (Supplementary Fig. 3a). Alphafold2 predicts that ProP folds into a ClbP-like structure with conserved catalytic residues (Supplementary Fig. 3b). Given the promiscuity of several prodrug peptidases^16^, we speculated that ProP may be responsible for the zwittermicin produced by the Δ*zmaM* strain. While deleting the *proP* gene in UW85 (Δ*proP*) had no impact on zwittermicin production, deleting *proP* in the Δ*zmaM* background (Δ*Z*Δ*P*) led to a greater accumulation of prezwittermicin and a further reduction in zwittermicin (Supplementary Fig. 2). Thus, ProP is not essential for zwittermicin production but can activate prezwittermicin.

That we observe prezwittermicin accumulation both intracellularly and extracellularly in Δ*zmaM* and Δ*Z*Δ*P* cultures suggests an alternative release pathway independent of ZmaM. We induce zwittermicin production by promoting sporulation^22^, during which release of the mature endospore occurs through cell lysis^26^, and we speculate that this lysis leads to the release of cytoplasmic prezwittermicin. However, we cannot rule out the possibility that membrane transporters other than ZmaM can also export prezwittermicin (Fig. 1c). Overall, our observation that intracellular prezwittermicin accumulates in the Δ*Z*Δ*P* strain strongly supports the conclusion that ZmaM is capable of both exporting and activating zwittermicin.

As a final test of the requirement for ZmaM for secretion and activation, we sought to complement the Δ*Z*Δ*P* strain in *trans*. We inserted the *zmaM* gene under the control of the *zmaA* promoter in a non-essential locus in the *B. cereus* genome (in the *pulA* gene encoding a glycosidase) in the Δ*Z*Δ*P* strain (Supplementary Fig. 1c). The ectopic *zmaM* gene greatly decreased both intracellular and extracellular prezwittermicin accumulation and restored extracellular zwittermicin accumulation (Fig. 1c). These results further support our conclusion that ZmaM can export prezwittermicin and activate it into zwittermicin.

### ZmaM activates prezwittermicin isolated from *ΔZΔP in vitro*

We developed a protocol to isolate prezwittermicin, a molecule that would otherwise only be accessible through a complex total synthesis, from Δ*Z*Δ*P* culture supernatant. Several steps of reverse-phase chromatography yielded a sample of prezwittermicin which was composed of two isomers (Supplementary Fig. 4a). The identity of prezwittermicin was confirmed by incubating it with purified samples of WT and peptidase-deficient ZmaM containing an S89A mutation at the catalytic serine (Fig. 1d). Both isomers of prezwittermicin were hydrolyzed within an hour of incubation into zwittermicin by WT, but not S89A. Because the produced zwittermicin showed no peak splitting, we speculate that prezwittermicin is produced with an additional acyl chain beyond the previously hypothesized single linear lauroyl^15^. Based on MS^2^ fragmentation (Supplementary Fig. 4b), the isomers have similar daughter ions; if the isomeric difference existed in the non-acyl chain structure of prezwittermicin, the difference would be apparent in this fragmentation. This supports that prezwittermicin isomers differ in acyl chains similar to preedeine, prexenocoumacin, preamicoumacin and precolibactin, which are all produced with varying acyl chains^3,7,15,27^. Overall, we show that we can isolate prezwittermicin from our Δ*Z*Δ*P* strain and that ZmaM can activate both prezwittermicin isomers *in vivo* and *in vitro*.

### The ZmaM peptidase active sites face the transporter’s outer vestibule

We next determined the structure of ZmaM containing an E965Q mutation at the Walker B motif to impair ATP hydrolysis. We purified detergent-solubilized C-terminally 10xHis-tagged ZmaM E965Q in the presence of ATP (Supplementary Fig. 5). Using cryo-EM, we obtained a C2-symmetric density map of 2.75 Å overall resolution (Fig. 2a, Extended Data Table 1, and Extended Data Figures 1 and 2), from which we resolved the molecular architecture of homodimeric ZmaM (Fig. 2b). The two half-transporter modules dimerize into a complete type IV ABC transporter in an outward-open conformation, with domain-swapped bundles composed of transmembrane helices (TMs) 3, 4, 5, and 6 from one subunit and 1 and 2 from the other. As expected, the TM4-5 intracellular coupling helix of one subunit contacts the nucleotide binding domain (NBD) of the other subunit. Consistent with an outward-open conformation, the two NBDs form a closed dimer with clear density for one ATP and one Mg^2+^ ion per subunit bound at the NBD interface (Fig. 2c). The peptidase modules straddle the ABC transporter TMD, with their 3-helix TMDs spanning each side of the outer vestibule opening and TM2’ of the peptidase interacting extensively with TM2 of the transporter. The peptidase-transporter TM-TM contacts span 564 Å^2^ of buried surface area (BSA) on each ZmaM subunit, and the intracellular intermodule linker is disordered. The two hydrolase domains sit in close apposition over the transporter, with their active sites facing the open outer vestibule.

**Figure 2.**
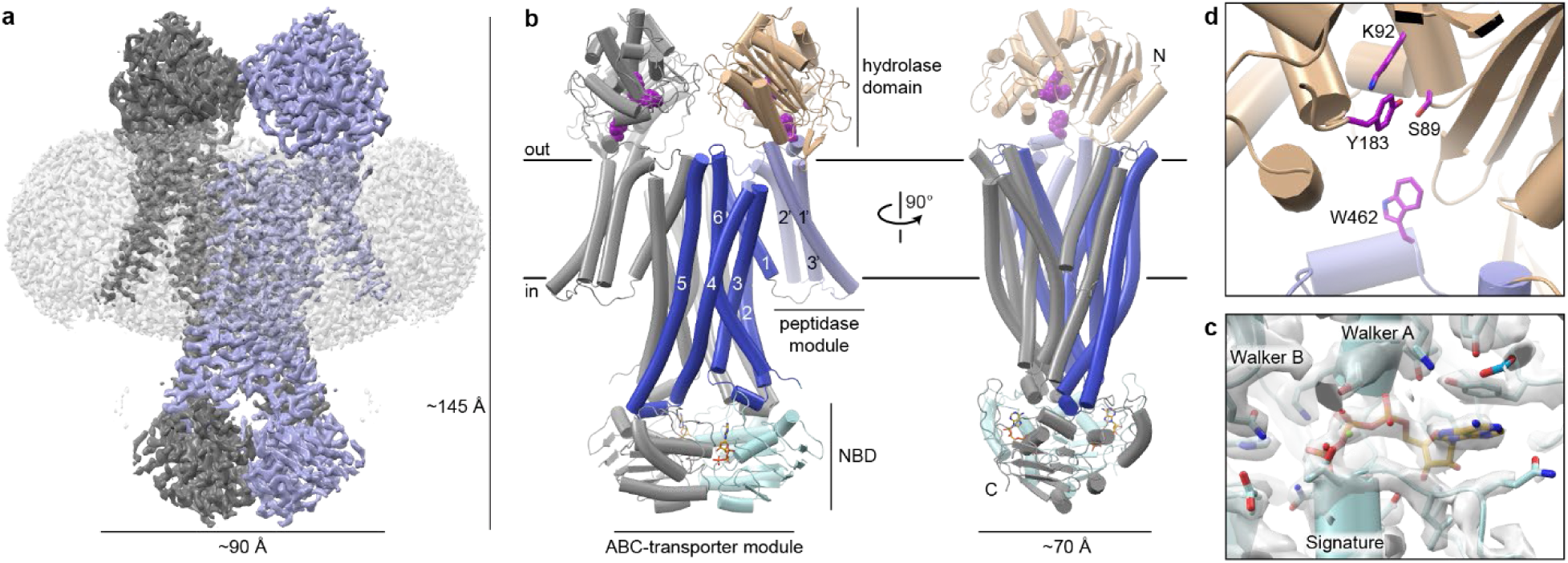
Structure of ZmaM. **a**, Cryo-EM map of ZmaM at 2.8-Å resolution contoured at 8σ. The maps of the two ZmaM subunits are colored grey and blue, and the micelle is shown as a transparent map contoured at 3σ. Overall dimensions of the ZmaM dimer are included. **b,** Cartoon representations of ZmaM. One subunit is grey and the other is colored by domain: the hydrolase and TMD of the peptidase module in wheat and light blue, respectively, and the TMD and NBD of the ABC transporter module in slate and cyan. The peptidase active site catalytic triads are highlighted as purple spheres. The view on the right is rotated by 90° with the grey peptidase module removed to show the open outer vestibule of the transporter. **c,** Density map of the ATP•Mg^2+^ bound at the NBD interface. Conserved NBD motifs are highlighted. **d,** Peptidase active site with conserved catalytic-site and substrate-binding sidechains highlighted as magenta sticks.

### The peptidase module TMD interacts extensively with the ABC transporter module

Many ABC transporters contain accessory TMDs that facilitate cellular localization and mediate interactions with partner proteins^28^. We sought to compare the position of the ZmaM peptidase modules to those of accessory domains in other type IV ABC transporters. We extracted available experimental structures of accessorized type IV ABC transporters from the InterPro database^29^ and retained structures of each paralog in each available conformation, yielding 9 structures of ABCB and ABCC family members (Supplementary Table 1). The ZmaM peptidase TMD interacts more extensively with the ABC transporter TMD than most other accessory TMDs (Extended Data Figure 3). In general, ABCC accessory domains have little to no TM-TM contacts with the transporter; instead, their position is fixed via extracellular loop contacts and a structured accessory domain-transporter linker. The human ABCC3 structure is an exception, with a total BSA of 885 Å^2^ between its accessory TMD and the transporter TMD, although it is unclear why its structure differs from the other ABCC homologs. The unusually substantial interface between the ZmaM peptidase and transporter TMDs suggests close coordination between their biochemical activities.

### The ZmaM peptidase module is similar to ClbP

The ZmaM peptidase modules resemble the structure of the type I prodrug peptidase ClbP (PDB ID:7UL6; RMSD values of 2.2 Å over 327 hydrolase domain Cα atoms, 2.7 Å over 52 TMD Cα atoms, and 5.6 Å overall over 408 Cα atoms), with the largest differences in the intracellular portions of TM1’ and TM2’ (Extended Data Figure 4a). ZmaM also shares the conserved active site residues—S89, K92, Y183, and W462—characteristic of all *N*-acyl-ᴅ-asparagine prodrug peptidases (Fig 2d). Both active sites are under the canopy formed by the hydrolase domains above the outer vestibule of the transporter. Although the path is not completely sealed (Fig. 2b), egress of the transported prezwittermicin substrates proximal to the peptidase active sites likely promotes efficient zwittermicin production.

The ZmaM peptidase module dimer is also similar to the ClbP dimer. The ZmaM hydrolase domains face each other like in ClbP (Fig. 3a), although the two TMDs are each splayed out 15° farther to accommodate the transporter module. Consistent with the lack of sequence conservation of the hydrolase domain dimerization interface^4,18^, the ZmaM and ClbP interfaces are very different. The ZmaM interface is minimal and lacks ClbP’s interlocking loops, with a BSA of 468 Å^2^ per subunit compared to 1296 Å^2^ in ClbP (Fig. 3b,c). This suggests that ZmaM dimerization is driven largely by the ABC transporter module. To test this hypothesis, we purified the isolated ZmaM peptidase module (hereafter ZmaM-PEPM; 58.1 kDa) and compared its size exclusion chromatogram with ClbP (55.4 kDa monomer; Fig. 3d). The larger retention volume for ZmaM-PEPM compared to ClbP, which we previously determined is dimeric in solution^18^, suggests that ZmaM-PEPM predominantly exists as a monomer in solution.

**Figure 3.**
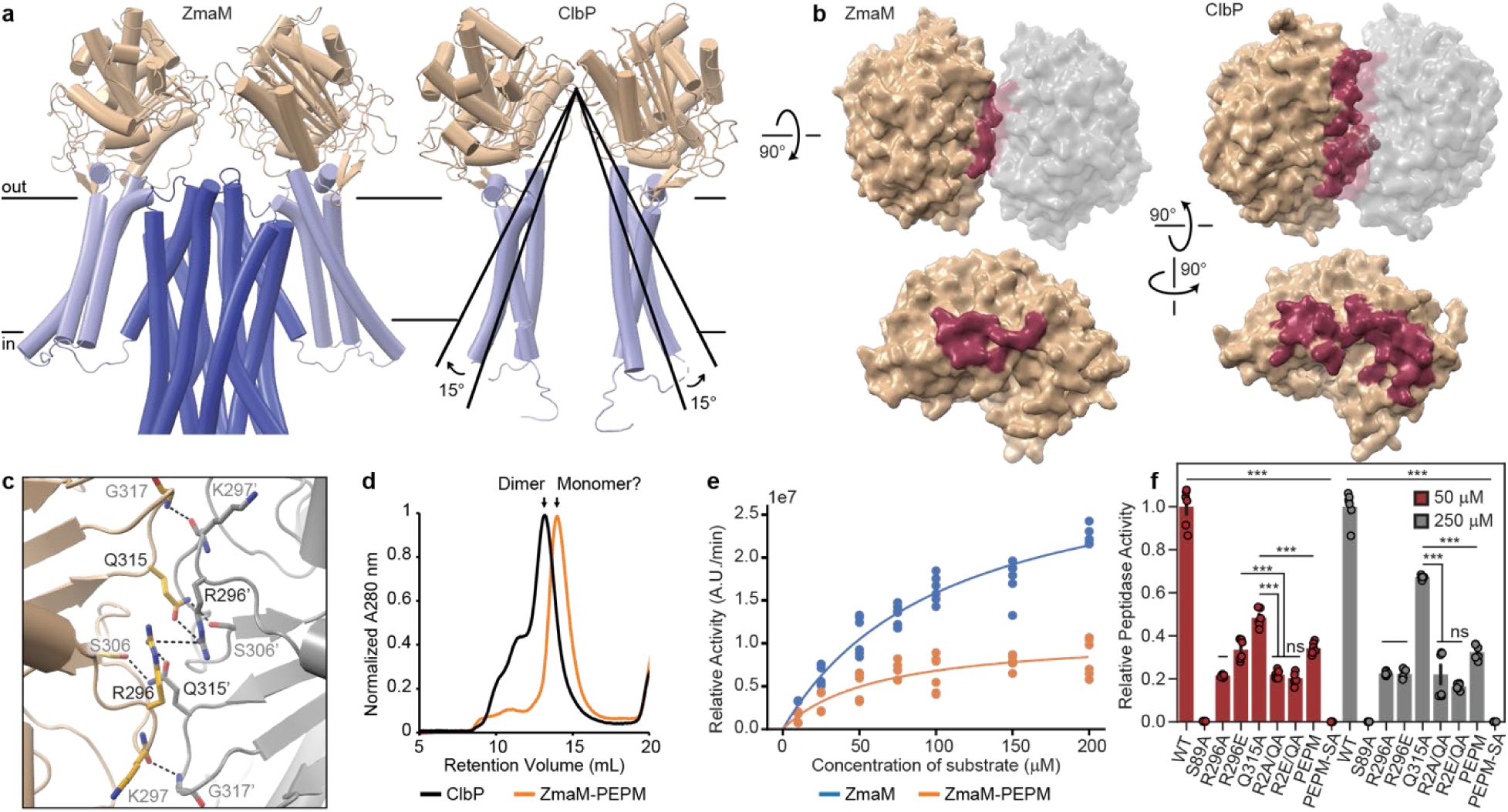
Features of the ZmaM peptidase module and comparisons to the type I prodrug peptidase ClbP. **a**, Side-by-side comparison of the ZmaM peptidase module with ClbP (PDB ID: 7MDE). **b,** Top view of the hydrolase domain dimer (top) and head-on view of the interface on one subunit (bottom) for ZmaM (left) and ClbP (right). BSA is highlighted in red on the surface representation. **c,** Hydrolase domain interface of ZmaM, with residues within 4 Å labeled and polar interactions highlighted with dotted lines. **d,** Size exclusion chromatograms for ClbP and ZmaM-PEPM (n = 2 biological replicates with 2 technical replicates each). **e,** Dose-response and Michaelis-Menten fits for ZmaM and ZmaM-PEPM *in vitro* peptidase activity (n = 6 replicates). **f,** Relative in vitro peptidase activities of ZmaM variants with hydrolase interface mutations and ZmaM-PEPM, normalized to WT (n = 6 replicates) at 50 μM (red) or 250 μM (gray) fluorogenic substrate concentration. P-values were obtained from *t*-tests and adjusted for multiple comparisons using a false discovery rate correction (not significant (ns), FDR-adjusted p > 0.05; *, p < 0.05; **, p < 0.01; ***, p < 0.001).

### ZmaM-PEPM is less active *in vitro* than full-length ZmaM

We sought to determine whether the ZmaM-PEPM peptidase activity is influenced by its assembly into the ZmaM dimer. We used a fluorogenic assay^17^ to measure the *in vitro* peptidase activity of purified ZmaM and ZmaM-PEPM. The dose-response curves reveal marked differences in Michaelis-Menten kinetic parameters (Fig. 3e). ZmaM-PEPM has a higher apparent affinity than full-length ZmaM (K_M_ = 110 ± 15 vs. 60 ± 17 μM for ZmaM and ZmaM-PEPM, respectively) but a lower V_max_ (32.3 ± 2.3 vs. 11.1 ± 1.2 A.U./min). To determine whether these differences arise from specific structural interactions at the hydrolase dimer interface, we introduced the mutations R296A, R296, and Q315A, as well as the R296A-Q315A (RQAA) and R296E-Q315A (RQEA) combinations (Fig. 3c). All variants had impaired activity, with the double-mutations variants least active (Fig. 3f). The similar effects on activity at two different substrate concentrations are consistent with most mutations primarily affecting V_max_ rather than K_M_. These results suggest that the inter-hydrolase interactions are important for peptidase activity in ZmaM, either for overall stabilization of the protein or for some allosteric effect influencing peptidase activity.

To analyze sequence conservation at the type II peptidase interfaces, we used a curated sequence list of prodrug peptidases^18^ along with our ZmaM sequence. Indeed, both R296 and Q315 are quite conserved in type II peptidases (R296 – 74%, Q315 – 73% conserved), whereas analogous positions are not conserved in type I peptidases (Extended Data Figure 4b). This suggests that in most type II prodrug peptidases R296 and Q315 promote interactions between the hydrolase domains that are important to the integrity of peptidase activity.

### ZmaM ABC transporter module conformations

Our ZmaM structure reflects an outward-open ABC transporter conformation (Fig. 2b). We used Colabfold^30–32^ and the AlphaFold3 webserver^33^ to predict additional conformations of ZmaM. Colabfold reliably predicted an inward-open conformation while AlphaFold3, when including two ATP ligands, predicted an occluded conformation (Fig. 4a). We used DALI^34^ to identify their closest structural neighbors. The inward-open model was most similar to the inward-open conformation of TmrAB^35^ (RMSD of 2.9 Å over 508 residues; PDB ID: 6RAG), the occluded model to the outward-occluded conformation of MsbA^36^ (RMSD of 2.7 Å over 511 residues; PDB ID: 7PH4) and our experimental structure to the outward-open conformation of TmrAB^37^ (RMSD of 2.6 Å over 489 residues; PDB ID: 6QUZ). These DALI results are consistent with the assumption that ZmaM follows a typical type IV ABC transporter conformational cycle and suggest that the AlphaFold models represent potential conformations of ZmaM.

**Figure 4.**
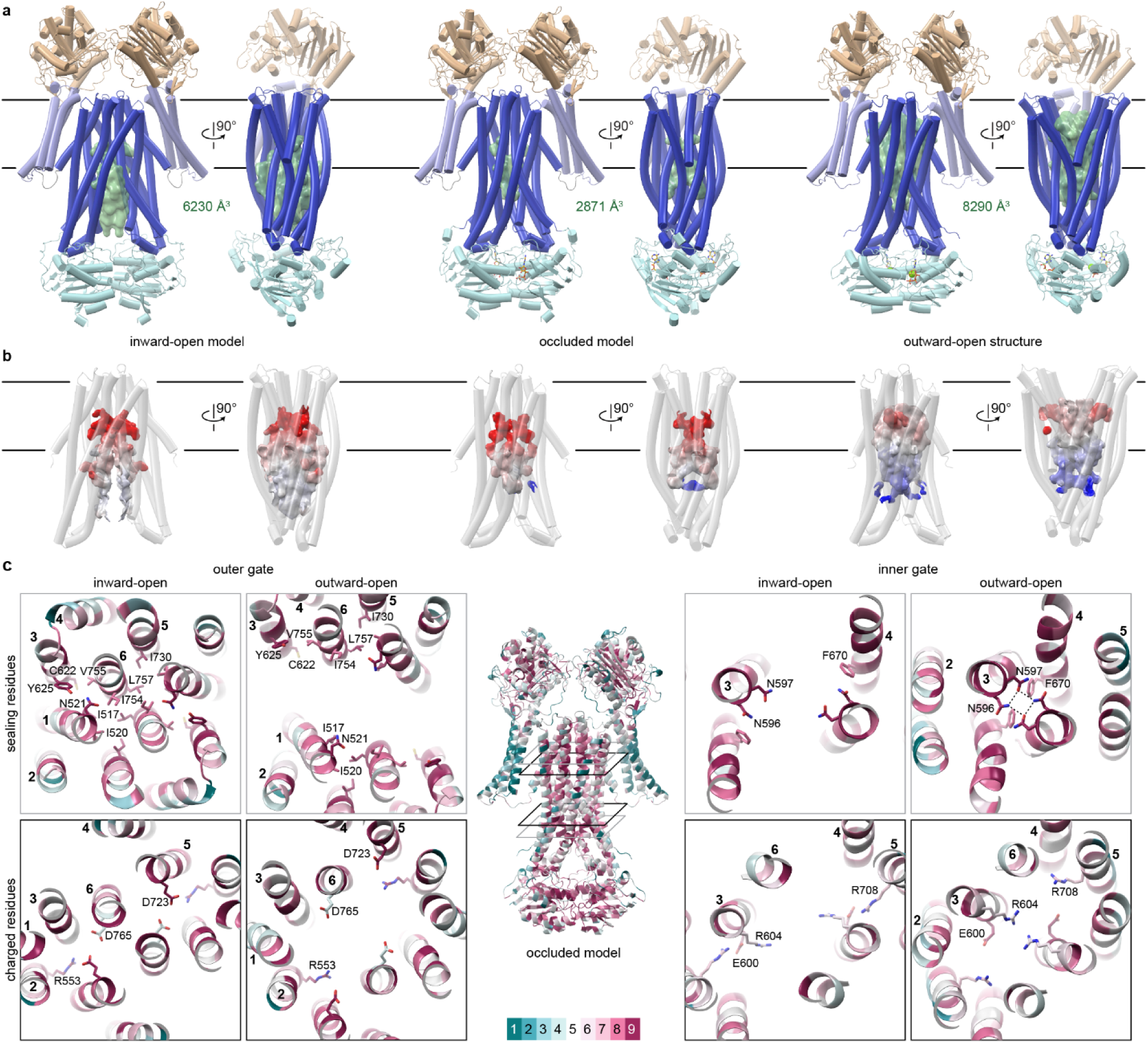
Predicted alternate conformations of ZmaM and their hints at transport details. **a**, Inward-open model, occluded model, and outward-open structure of ZmaM including head-on (left) and side views (right; front peptidase module omitted to better visualize the transporter module). Predicted transporter cavities are displayed as green surfaces, and their volumes indicated. **b,** Transporter module TMD as transparent cartoon with surface representation of the cavities colored according to electrostatics as predicted by APBS^66^, from -20 kcal/mol (red) to +20 kcal/mol (blue). **c**, All panels are colored by ConSurf^63^ score of sequence conservation across type II peptidases from cyan (variable) to magenta (conserved). All panels are views from the extracellular side, and their positions are indicated on the center model. Both the outer (left) and inner (right) gates have highly conserved sealing residues (top, grey-lined panels). Less conserved charged residues line the substrate cavity proximal to each gate (bottom, black-lined panels), with primarily negatively charged residues at the outer gate and positively charged residues at the inner gate.

The presence of two hydrolase active sites per transporter raises the question of whether ZmaM naturally processes one or two prezwittermicins in each transport cycle. Prezwittermicin has a predicted volume of 681 Å^3^, whereas the ZmaM transporter cavity volume is >6230, 2871, and >8290 Å^3^ for the inward-open, occluded, and outward-open conformations, respectively (Fig. 4a). Thus, the predicted inward-open conformation may represent the substrate binding-ready state, and our outward-open structure the post-release structure. Furthermore, the occluded cavity can readily accommodate prezwittermicin, and perhaps more than one, as has been observed in some ABC transporters^38,39^.

Prezwittermicin is predicted to be positively charged (Fig. 1b). On the electrostatic potential surface of the inward-open ZmaM model (Supplementary Figure 6a), the deepest point of the predicted cavity has a negative potential arising from two pairs of aspartates (Fig. 4b-c), which could attract and stabilize bound prezwittermicin. The occluded cavity is analogously negative at its extracellular end, but more positive at its intracellular end. This positive surface, arising from two pairs of arginines (Fig. 4c), remains in the outward-open structure and may promote prezwittermicin release. In contrast, the substrate-binding sites of the hydrolase domains have a negative surface (Supplementary Figure 6a), which may help attract the substrate. This analysis suggests that surface charges may guide prezwittermicin throughout its transport cycle and activation.

These charged residues lining the ends of the substrate-binding cavity are poorly conserved across type II prodrug peptidases (Fig. 4c), as are many of the cavity-lining residues (Supplementary Figure 6b-d), consistent with the distinct peptidases being adapted to their specific prodrug substrates. In contrast, the residues forming the outer- and inner-gate seals are highly conserved (Fig. 4c), consistent with their conserved function facilitating the transporter conformational cycle.

### Isolated transporter module is less active than in full-length ZmaM

Given that ZmaM-PEPM is less active than full-length ZmaM, we sought to determine whether isolating the ABC transporter module (ZmaM-ABCM) also affects its activity. We first reintroduced the coding sequence for either ZmaM-PEPM or ZmaM-ABCM into the *B. cereus* Δ*Z*Δ*P* strain. While reintroducing ZmaM-PEPM did partially rescue the production of zwittermicin, reintroducing ZmaM-ABCM did not increase the amount of secreted prezwittermicin (Supplementary Fig. 7). This suggests that ZmaM-ABCM is either poorly expressed or inactive. We also measured the *in vitro* basal ATPase activity of purified ZmaM and ZmaM-ABCM as a proxy for basic transporter function. While ZmaM hydrolyzed ATP with a K_M_ of 1.35 ± 0.37 mM consistent with values reported for other ABC transporters^40–42^, ZmaM-ABCM showed virtually no ATPase activity *in vitro* (Fig. 5a). Overall, our results suggest that either through allostery or stability, the presence of the peptidase module is important for proper function of the ZmaM ABC transporter module.

**Figure 5.**
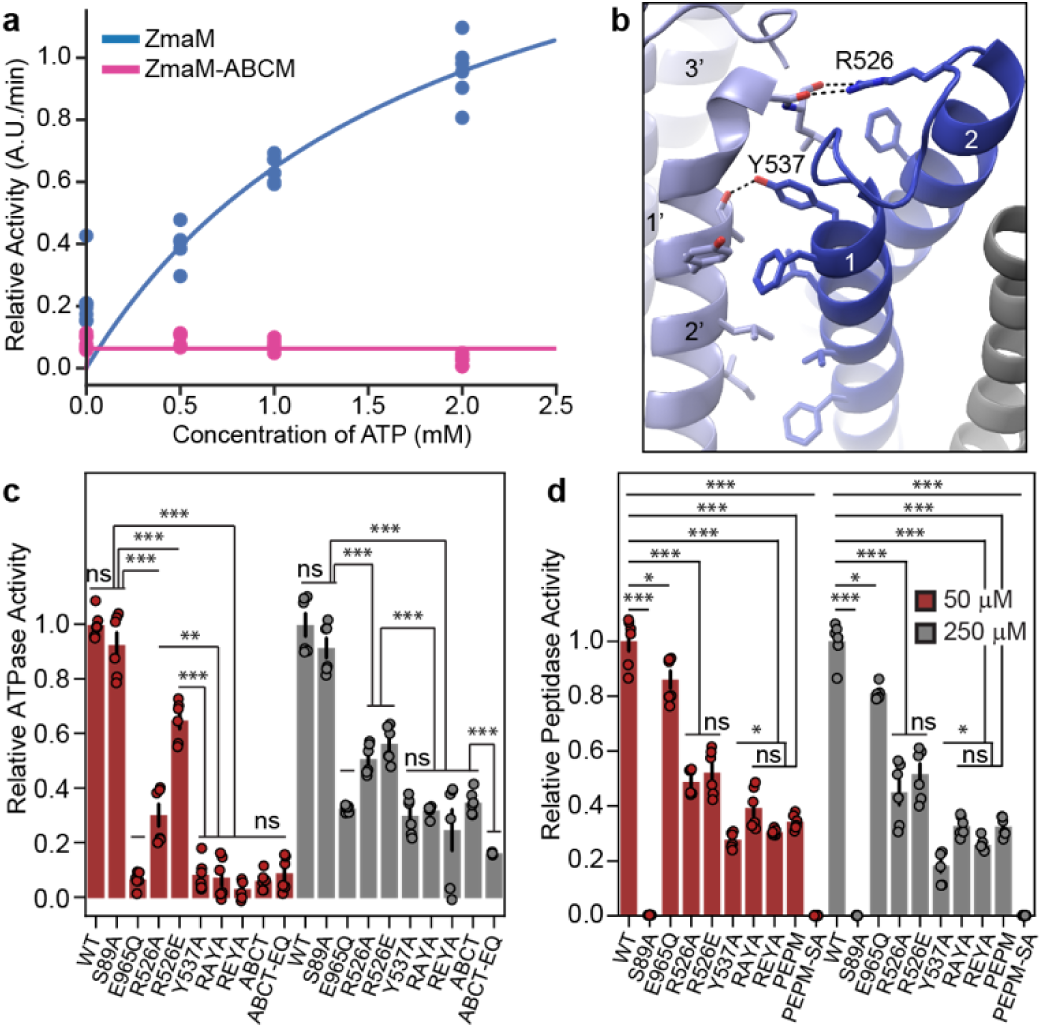
Specific sites may hint at inter-module dependence. **a**, Dose response and Michaelis-Menten fits for ZmaM and ZmaM-ABCM basal ATPase activity (n = 6 replicates). **b,** The TMD interface between the two ZmaM modules, with residues within 4 Å shown in sticks and mutated residues—ABC transporter residues R526 and Y537 interacting with main chain carbonyls from ZmaM-PEPM—labeled. **c,** Relative *in vitro* basal ATPase activities of ZmaM, ZmaM variants with inter-module interface mutations, and ZmaM-ABCM, normalized to WT (n = 6 replicates) at 1 mM (red) or 5 mM (gray) ATP. **d,** Relative *in vitro* peptidase activities of ZmaM variants with inter-module interface mutations and ZmaM-PEPM, normalized to WT (n = 6 replicates) at 50 μM (red) or 250 μM (gray) fluorogenic substrate concentration. For both **c,** and **d,** p-values were obtained from *t*-tests and adjusted for multiple comparisons using a false discovery rate correction (not significant (ns), FDR-adjusted p > 0.05; *, p < 0.05; **, p < 0.01; ***, p < 0.001).

### Inter-module contacts are important for ZmaM activity

To further investigate this inter-module dependency for ATPase and peptidase activities (Fig. 3e and 5a), we sought to identify interface residues that may be involved. In our outward-open structure, the ZmaM-PEPM TM2’ C terminus and ZmaM-ABCM TM1 interact (Fig. 5b). Although most of the interface is nonpolar, R526 interacts with backbone carbonyls from the TM2’ C terminus and Y537 interacts with the A442 carbonyl left unpaired due to a kink in TM2’. We generated the R526A, R526E, and Y537A mutations, as well as the R526A-Y537A and R526E-Y537A double mutations. All variants have impaired ATPase activity, and only R526A and R526E have ATPase activity above ZmaM-ABCM (Fig. 5c). Consistent with the effect of the mutations, R526 is conserved in 40% and Y537 in 80% of the type II prodrug peptidases sequences (Extended Data Figure 4b). We also observed impaired peptidase activity across all these variants, and with their activity similar to ZmaM-PEPM’s activity (Fig. 5d). Collectively, these results indicate that the integrity of the inter-module interface is important for optimal activity of both ZmaM modules.

## Discussion

We chose the *B. cereus* UW85 ZmaM as a model type II prodrug peptidase. We demonstrated *in vivo,* by deleting and then *trans*-complementing the *zmaM* gene, that ZmaM is responsible for the final two steps of zwittermicin production, export and activation through hydrolysis of *N*-acyl-ᴅ-asparagine. The Δ*Z*Δ*P* strain allowed us to isolate the precursor prezwittermicin. With this isolated metabolite, we showed that ZmaM activates prezwittermicin *in vitro* as well. The cryo-EM structure of ZmaM in an outward-open conformation reveals close spatial proximity between the transporter vestibule and the peptidase active sites. This structure also showed that the peptidase modules dimerize in a manner akin to the type I peptidase ClbP. In ZmaM this dimerization is likely driven by the ABC transporter dimer because the isolated peptidase module was a monomer. We used Alphafold to predict alternate conformations for ZmaM. These models with the transporter module in inward-open or occluded conformations have substrate-binding cavities with surfaces conserved across type II prodrug peptidases and electrostatics consistent with prezwittermicin transport. Finally, we find interdependence between the modules of ZmaM, as both modules are less active when isolated or when inter-module interface residues are mutated.

Of note, our *B. cereus* genetic engineering also uncovered alternative pathways for prezwittermicin export and activation. In the absence of ZmaM, prezwittermicin release is likely facilitated by cell lysis during sporulation, albeit this is non-specific. We also identified an orphan *N*-acyl-ᴅ-asparagine prodrug peptidase, ProP, which produced some zwittermicin in the absence of ZmaM, indicating that its expression overlaps at least partly with ZmaM. Its genomic neighborhood does not include NRPS or PKS enzymes typically associated with *N*-acyl-ᴅ-asparagine prodrug toxins. The physiologically relevant substrate(s) of ProP therefore remain unclear.

When isolating prezwittermicin from the *zmaM* knockout strains, we found two forms with slightly different elution times, and our purified prezwittermicin similarly contains both forms (Fig. 1c,d). Consistent with zwittermicin showing a single elution peak, we found evidence that the two prezwittermicin forms differ in their prodrug motif acyl chain, akin to the different acyl substituents found in other *N*-acyl-ᴅ-asparagine prodrug toxins^3,7,15,27^. Isolated prezwittermicin will be a useful chemical tool to further investigate ZmaM’s function and biochemical mechanisms.

ZmaM is a member of the accessorized ABC transporters, and we have evidence that its two modules are interdependent, as full enzymatic activity requires both modules and intact inter-module interfaces. It is still unclear whether this interdependence is the result of structural stability provided by the modular assembly, or whether the transport and peptidase activity are allosterically regulated through inter-module interactions. In PCAT-type dual peptidase-transporters, peptidase activity is regulated by the transporter’s conformational state^43^. This ensures that the toxin is activated only when it can be immediately transported, protecting the producing bacterium. However, type II prodrug peptidases do not require analogous allostery as toxin activation occurs extracellularly, although allosteric coordination of transport and peptidase activity could maximize active toxin production.

The ZmaM peptidase module dimer is reminiscent of ClbP’s dimer, although the hydrolase domain interfaces are completely different. Because of the lack of sequence conservation, we previously proposed that the ClbP dimer may have evolved to optimize activation of pseudodimeric precolibactin^18^, whereas zwittermicin and other known prodrug toxins only contains one prodrug motif. In both ClbP^18^ and ZmaM, disrupting the interface between the two hydrolase domains impairs peptidase activity. The similarities between the ClbP and ZmaM dimers raise the question of whether some type I peptidases may also interact, although intermolecularly, with transporters in their own pathways. ClbP has been hypothesized to interact with ClbM^18^, a multidrug and toxic compound extrusion (MATE) transporter that exports precolibactin^44^. ClbP also plays an essential but poorly understood role in microcin production by *E. coli* Nissle 1917. This role requires only the ClbP TMD, raising the possibility that it interacts with other components of the microcin biosynthetic machinery such as the ABC exporter MchF^45^.

The structural information on ZmaM and new tools like the Δ*Z*Δ*P* strain and isolated prezwittermicin will facilitate further studies of ZmaM as a model type II prodrug peptidase, and of its homologs. Elucidating the determinants of substrate specificity or promiscuity in their transporter and peptidase modules will be important for regulating or engineering their activities for therapeutics or biotechnology applications. Our new mechanistic understanding of ZmaM represents an important first step towards identifying the molecular targets of type II prodrug peptidases and developing therapeutics and biotechnological tools based on these systems.

## Materials and Methods

### ZmaM plasmid construction

ZmaM was amplified by PCR from *B. cereus* UW85 genomic DNA (provided by the Balskus lab) and inserted into pET21-C10H (derived from pET21a by inserting the 5’-TCATCATCACCATCACCACCATCACCATCATTGAGGCCGCA-3’ sequence between its NotI and XhoI sites) at the NdeI and NotI sites, yielding the construct used for structure determination with a C-terminal 3xAla linker and 10xHis tag. For *in vitro* assays, the construct was modified to minimize the co-purification of an N-terminally truncated product produced from an alternative translation start site (see Supplementary Note 1). Briefly, we mutated a non-canonical start site at the V742 GTG codon to GT<u>T</u> (synonymous mutation), introduced the conservative M744L mutation (ATG to <u>C</u>TG) and synonymous mutations to a possible internal Shine-Dalgarno sequence (GGAGAT to <u>A</u>GA<u>A</u>AT mutations), we introduced an N-terminal 8xHis-tag after the signal peptide and removed the C-terminal 10xHis-tag by using 3-piece Gibson Assembly (Gibson Assembly Master Mix, NEB E2611S). The coding sequence from the signal peptide through residue 498 of this N-terminally 8xHis-tagged construct was inserted between the NdeI and NotI sites of pET21a to generate the ZmaM-PEPM construct. The ZmaM-ABCM coding sequence consists of residues 488-1027 cloned into pET21-N8H^46^. All point mutations were introduced using site-directed mutagenesis and verified by full plasmid sequencing (Plasmidaurus). Supplementary Table 2 lists primers used for all cloning.

### *B. cereus* plasmid construction

Plasmids to knock out genes in *B. cereus* were constructed using a Rudner lab pMAD-derived vector, pFR50. This vector carries a thermosensitive origin of replication, an erythromycin resistance cassette for *Bacilli*, and mCherry fused to the *B. subtilis veg* promoter (Supplementary Fig. 1b, c). Allelic exchange plasmids containing ∼1000-bp upstream and downstream of the gene to be knocked out were assembled using Gibson Assembly. The loop-out retained 3 N-terminal and 7 C-terminal codons to minimize polar effects. For *zmaM trans* complementation, the *zmaM* ORF with 300-bp of the upstream *zmaA* promoter region was inserted into a plasmid harboring front and back halves of the non-essential *pulA* gene and an antibiotic resistance cassette, analogous to the commonly used *amyE* locus in *B. subtilis*^25^. The flanking homology regions were each 600 bp (Supplementary Fig. 1c). Plasmids were prepared from *dam-/dcm-E. coli* cells (New England Biolabs, C2925H) before transformation into *B. cereus*. Supplementary Table 2 lists primers used for these constructs.

### *B. cereus* genetic knockouts and *trans* complementation

*B. cereus* electrocompetent cells were prepared as follows: one freshly streaked *B. cereus* colony was cultured in lysogeny broth (LB) at 37°C to an OD_600_ of ∼ 0.4 – 0.5, then diluted 1:50 into pre-warmed LB supplemented with 200 mM sucrose and 0.1 µg/mL tunicamycin and grown to OD_600_ of ∼ 0.3. After 3 minutes on ice, cells were pelleted at 4,000 × g at 4°C for 10 minutes, washed twice with 0.2 and twice with 0.02 volume ice-cold electroporation buffer (0.5 M sucrose, 10% glycerol), then resuspended in 0.004 volume of electroporation buffer.

Unmethylated plasmid (0.5-1 µg in 10 µL) was transformed in 100 µL of cells by electroporation in 1-mm cuvettes using a Bio-Rad Gene Pulser II (25 µF, 200 Ω, 18 kV/cm). Pre-warmed LB supplemented with 20 mM MgCl_2_ and 2 mM CaCl_2_ (900 µl) was added and the cells allowed to recover at 37°C for 2 hours before plating on LB supplemented with erythromycin (5 µg/mL; LB-Erm) and overnight incubation at 30°C.

To initiate the loop-in-loop-out process, transformants were grown in 5 mL LB-Erm at 30°C for 6-8 hours, then 100 µL of serial dilutions (10^-1^, 10^-3^, 10^-5^) were plated on LB-Erm and incubated at 42°C for 16-24 hours to select for integrated plasmids (loop in). A single integrated colony was then grown without selection in 5 mL LB at 30°C for 6-8 hours to promote loop-out. Serial dilutions (100 µL at 10^-5^, 10^-6^, 10^-7^) of the saturated culture were plated on LB and incubated at 30°C for 16 hours. Colonies were screened for sensitivity to erythromycin, and allelic exchange was confirmed by colony PCR and amplicon sequencing (Plasmidsaurus). Confirmed strains were frozen as glycerol stocks.

### ZmaM purification for cryo-EM

C-terminally 10xHis-tagged ZmaM was expressed in *E. coli* C41(DE3) (Lucigen). A single colony was grown in 120 mL LB overnight. The culture was diluted 1:100 into a total of 12 L Terrific Broth with 100 mg/mL ampicillin, grown at 37°C until OD_600_ of 0.8-1.0, and induced with 0.5 mM isopropyl β-D-1-thiogalactopyranoside for 20 hours at 15°C. Cells were harvested by centrifugation at 3635 × g for 20 minutes and either flash frozen or immediately prepared for lysis. All purification steps were performed at 4°C. The cell pellet was resuspended in lysis buffer (20 mM sodium phosphate pH 8, 500 mM NaCl, 55 mM imidazole, 1 mM benzamidine, 1 mM phenylmethane sulfonyl fluoride, 0.3 mg/mL DNAse, 0.3 mg/mL lysozyme) at a ratio of 2 mL/g of cell pellet and disrupted using a Branson Sonifier 450 with the following settings: 65% duty cycle, 10 output control, 45 seconds on, 45 seconds off for 3 cycles and 90 seconds off for an additional 3 cycles. Cell lysates were clarified by centrifuging at 27100 × g for 15 minutes, and membranes pelleted from the clarified lysate at 45000 × g for 75 minutes. Membranes were homogenized using a glass Potter–Elvehjem grinder into lysis buffer supplemented with 1 mM ATP and 5 mM MgCl_2_, and either flash frozen or immediately solubilized. Membranes were solubilized by adding 1% n-dodecyl-β-D-maltopyranoside (DDM, Anatrace) and rocking for 1 hour, centrifuged at 35000 × g for 35 minutes, then loaded on ∼2 mL pre-equilibrated Ni-NTA Sepharose resin (Qiagen), rocking for 1 hour. The resin was loaded into a gravity column, washed with 10 column volumes (CV) of wash buffer (lysis buffer with 1 mM ATP, 5 mM MgCl_2_, and 0.02% DDM), followed by step 3-CV elutions with imidazole increased to 75 mM, 100 mM, 150 mM, 250 mM, 300 mM, and 450 mM. Typically, the 250 mM and 300 mM eluates were pooled and concentrated using a 100-kDa MWCO concentrator (Millipore) to <500 µL before running onto a Superose 6 10/300 GL (Cytiva) size exclusion chromatography (SEC) column in 10 mM Tris pH 8, 150 mM NaCl, 1 mM DTT, and 0.02% DDM), collecting 250-µL fractions. The purest and most concentrated fractions were pooled and concentrated using a 100-kDa MWCO concentrator to 3.9 mg/mL, then flash frozen.

### Grid sample preparation and imaging

Quantifoil Cu R1.2/1.3 400 mesh grids were glow discharged using a PELCO easiGLOW Discharge System at 0.39 mBar, 15 mA for 30 seconds. Three µL of ZmaM sample were applied to the carbon side, then vitrified in 100% liquid ethane using a Vitrobot Mark IV (Thermo Fisher Scientific) with a wait time of 30 seconds, blot time of 3 seconds, and a blot force of 0 in 100% humidity at 22°C.

### Cryo-EM data collection and processing

Cryo-EM data were collected on a 300 kV Titan Krios G3i Microscope (Thermo Fisher Scientific) equipped with a K3 direct electron detector (Gatan) and a GIF quantum energy filter (20 eV) (Gatan) using counted mode at the Harvard Cryo-Electron Microscopy Center for Structural Biology at Harvard Medical School. Two separate datasets were collected for a total of 25,587 micrographs. Data were acquired using image shift and real-time comma correction by beamtilt using the automated data collection software SerialEM^47^; nine holes were visited per stage position acquiring two movies per hole. Details of the data collection and dataset parameters are summarized in Extended Data Table 1. Dose-fractionated images were gain-normalized, aligned, dose-weighted and summed using MotionCor2^48^. Contrast transfer function (CTF) and defocus value estimation were performed using CTFFIND4^49^. Details of the data processing strategy are shown in Extended Data Figure 1 and map evaluation metrics are shown in Extended Data Figure 2. In short, particle picking was carried out using crYOLO^50^ followed by Heterogeneous refinement in cryosparc^51^, to give 1,215,193 particles. Data from the two collections were subsequently subjected to Relion 3D refinement, CTF refinement and Bayesian polishing before combining the particle stacks. The combined particle stack was then subjected to Focused 3D classification without alignment using a mask enclosing the NBDs using six classes. Two of the classes had well resolved features for the NBDs. These were selected and subjected to Relion 3D refinement with blush regularization to yield a final reconstruction at 2.75 Å with C2 Symmetry imposed (2.84 Å C1). Structural biology applications used in this project were compiled and configured by SBGrid^52^.

### Model Building and Refinement

The starting model was created *ab intio* using ModelAngelo^53^, then subjected to whole model simulation in the ChimeraX plugin, ISOLDE^54^. The resulting model was further refined with PHENX real space refinement^55^ using morph model, xyz minimization, local grid search, nqh flips, and adp refinement with Ramchandran and secondary structure restraints. After iterations of Coot^56^ rebuilding and fixing of outliers using Isolde, the model was further refined in PHENIX using morph model, xyz minimization, adp refinement with reference model restraints using the input model as the reference, Ramachandran and secondary structure restraints. Minor adjustments to some rotamers were made based on predicted hydrogen bonds between residues in the inter-hydrolase interface, and model finalized after a last round of adp refinement.

### Protein purification for activity assays

ZmaM-PEPM and ABCT constructs were expressed similarly to full-length ZmaM, except the cultures were transferred to 15°C at an OD_600_ of 0.3 before inducing at OD_600_ of 0.6. N-terminally 8xHis tagged full-length ZmaM and its variants were expressed as described above. All proteins used for activity assays were purified as described above, except that the SEC was replaced with a buffer exchange using PD-10 desalting columns (Cytiva). Eluates were concentrated using 100-kDa MWCO concentrators to 0.5-1 mg/mL and flash frozen.

### Fluorogenic peptidase assay

Fluorogenic assay was performed as described^17^. Briefly, in a 384-well all-black square flat-bottom plate, reactions contained 50 nM protein and 50 μM fluorogenic probe in 25 μL of 50 mM Tris-HCl pH 8.0, 200 mM NaCl, 0.025% DDM, 1 mM ATP, and 5 mM MgCl_2_. The assay was initiated by adding a protein master mix to a probe master mix. Fluorescent signal from the hydroxycoumarin (excitation: 340 nm, emission: 442 nm) was tracked using a SpectraMax i3 plate reader (Molecular Devices). Initial velocities were estimated from the slope of fluorescent signal generated from a linear fit of the data collected from 20-40 minutes. Triplicates were collected on the same plate, and relative activity was determined with respect to the triplicate average of WT ZmaM.

### ATPase activity assay

ATPase activity was determined using the Enzchek Phosphate Assay Kit (Invitrogen). In a 384-well clear square flat-bottom plate, reactions contained 350 nM protein, 1X assay reaction buffer (50 mM Tris-HCl pH 7.5, 1 mM MgCl_2_, 0.1 mM sodium azide), 0.25 mM 2-amino-6-mercapto-7-methylpurine riboside, 1.0 U purine nucleoside phosphorylase, and 5 mM ATP in final volume of 25 μL. Reactions were initiated by adding a protein and assay reaction master mix to an ATP master mix. Absorbance at 360 nm was tracked using a SpectraMax i3 plate reader (Molecular Devices). Initial velocities were estimated from the slope of fluorescent signal generated from a linear fit of the data collected from 20-40 minutes. Triplicates were collected on the same plate, and relative activity was determined with respect to the triplicate average of WT ZmaM.

### Prezwittermicin purification

A single Δ*Z*Δ*P* colony was inoculated into 4 L of 0.5X tryptic soy broth and grown at 30°C shaking at 200 rpm for 72 hours. The culture was filtered through a 0.2-micron nylon filter, then incubated with 80 g of HP-20 resin (Sigma-Aldrich) overnight at 4°C with light mixing. The following day, the resin was loaded into a column, the flowthrough was discarded, followed by elution with 100 mL of 100% methanol. The eluate was dried using a rotary evaporator (Labarota 4300, Heidolph) then resuspended in 40 mL of water and applied to a Sep-Pak C18 6 cc Vac Cartridge (Waters), washed with a stepwise gradient of 3 CVs each of 2%, 10%, 20%, and 40% methanol, and eluted with 3 CVs 100% methanol. This eluate was again dried using a rotary evaporator to yield a crude prezwittermicin powder (approx. 300 mg). The solid was dissolved in 3.5 mL of 25% acetonitrile / 75% water and filtered through a 0.2 µm 13 mm nylon filter (Pall). The sample was purified over three injections by preparatory HPLC (LCPrep AutoPure, Waters) on a 19x250 mm XBridge-C8 column (Waters) with the following gradient (30 mL/minute, collecting 15-mL fractions): 90% A (water + 0.1% formic acid) and 10% B (acetonitrile + 0.1% formic acid) for 5 minutes, gradient to 95% B for 10 minutes, hold at 95% B for 5 minutes, gradient to 10% B for 4 minutes, hold at 10% B for 6 minutes. Fractions containing prezwittermicin were pooled, concentrated under reduced pressure and lyophilized to dryness, yielding approximately 12.5 mg. The sample was resuspended in 1.6 mL of 35% acetonitrile / 65% water, filtered through a 0.2 µm 4 mm nylon filter (Pall), and repurified over three injections on the XBridge-C8 column with the following gradient (20 mL/minute, collecting 6.7-mL fractions during isocratic elution): 65% A and 35% B for 13 minutes, gradient to 95% B for 5 minutes, hold at 95% B for 4 minutes, gradient to 35% B for 2 minutes, hold at 35% B for 6 minutes. Fractions containing prezwittermicin were concentrated and lyophilized, yielding two fractions containing mostly the major (∼5.0 mg) and minor (∼1.5 mg) prezwittermicins. Each fraction was resuspended (major in 1.6 mL; minor in 1.0 mL) in 60% methanol / 40 % water, filtered through a 0.2 µm 4 mm nylon filter, and purified over two (major) or one (minor) injections on a 19x250mm Atlantis-T3 HPLC column (Waters) with the following gradient (20 mL/minute, collecting 3.3-mL fractions during isocratic elution): 40% A and 60% C (methanol + 0.1% formic acid) for 13 minutes, gradient to 95% C for 5 minutes, hold at 95% C for 4 minutes, gradient to 60% C for 2 minutes, hold at 60% C for 6 minutes. Fractions containing prezwittermicin were concentrated and lyophilized to afford the major (1.5 mg) or minor (0.2 mg) prezwittermicins.

### LC-MS analysis

Single *B. cereus* colonies were inoculated into 10 mL of 0.5X tryptic soy broth and shaken at 30°C at 200 rpm for 72 hours. Cells (intracellular samples) and supernatants (extracellular) were separated by centrifugation at 4000 × g for 20 minutes at 4°C. Supernatants were filtered through 0.2 micron PES syringe filters (VWR). Cell pellets were washed twice by resuspending in 5 mL phosphate-buffered saline and repelleting. Supernatant and cell pellet samples were stored at -80°C until further analysis. To prepare samples for LC-MS, supernatants (150 µL) were mixed with equal parts of acetonitrile (150 µL), then filtered through a 0.2 µm wwPTFE filter (Pall), and placed in autosampler vials. Pellets were extracted with 300 µL of extraction solvent (2:2:1 methanol:acetonitrile:water), sonicated in a water bath for 5 minutes, centrifuged at 3200 × g for five minutes, then filtered through a 0.2 µm wwPTFE filter (Pall), and placed in auto sampler vials. Samples were analyzed using a Horizon Vanquish UHPLC coupled to a Thermo Orbitrap IQ-X Tribrid mass spectrometer. A Kinetex C8 column (1.7 µm, 100 Å, 150 × 2.1 mm, Kinetex) was used for prezwittermicin, and an ACQUITY UPLC BEH amide column (1.7 µm, 150 × 2.1 mm, Waters) for zwittermicin, injecting 2 µL of sample, with a flow rate of 0.4 mL/minute using mobile phase A = 0.1% formic acid in water and B = 0.1% formic acid in acetonitrile. The LC conditions for prezwittermicin were: 10% B for 2 minutes, gradient to 90% B for 27 minutes, hold at 90% B for 3 minutes, then hold at 10% B for 3 minutes. The MS settings for prezwittermicin were: mass range 70–1200 m/z, 120000 resolution, the RF lens was 35%, the standard AGC target was used, and the auto maximum injection time mode was used. For MS/MS analysis, the ion of interest was acquired with a 1.6 *m/z* isolation window, using an ion trap with rapid scan rate, standard AGC target, and higher collision-induced dissociation (HCD) fragmentation with 30% collision energy. The LC conditions for zwittermicin were: 95 % B for 2 minutes, gradient to 40% B for 27 minutes, hold at 40% B for 3 minutes, fold at 95% B for 3 minutes. The MS settings for zwittermicin were: mass range 70–700 m/z, 240000 resolution, the RF lens was 35%, the standard AGC target was used, and the auto maximum injection time mode was used. Spectra were acquired in positive mode. A mass window of 2 ppm was used for extracted ion chromatograms.

For *in vitro* prezwittermicin hydrolysis assays, we incubated 50 nM of purified ZmaM construct with 50 μM prezwittermicin in 20 μL of 50 mM Tris pH 8.0, 200 mM NaCl, 1 mM ATP, 5 mM MgCl_2_, 0.02% DDM, and 2% DMSO at 25°C for 1 hour. The reaction was quenched with 180 μL acetonitrile and flash frozen. Samples were thawed and spun (10 min x 20,000 g) and processed for LC-MS as supernatant samples above.

### Structural analyses

The inward-open ZmaM conformation was predicted using ColabFold-multimer^30–32^ using default settings, two subunits predicted, and with templates on. The occluded conformation was predicted with the AlphaFold3 web server using default settings, two subunits predicted, and with two molecules of ATP included. Cavities were calculated using CavitOmiX (v. 1.0, 2022, Innophore GmbH). The cavities were calculated using a modified LIGSITE algorithm^57^. Surface electrostatics of all conformations were done with the Adaptive Poisson-Boltzmann Solver (APBS 3.4.1) plugin^58^ in PyMOL v.3.1.3.1. RMSD values were calculated using the align command in PyMOL; buried surface areas were calculated in PyMOL. The prezwittermicin volume was predicted using Molinspiration cheminformatics^59^.

### Sequence bioinformatics

To build the type II prodrug peptidase alignment, we extracted the relevant sequences from our published alignment^18^. Briefly, all sequences for the β-lactamase superfamily (PF00144) were downloaded from Uniprot and sequences between 400-1200 amino acids with at least 2 transmembrane helices were retained. These sequences were then further curated by a combination of scaffold annotation and genome neighborhood analysis to curate 271 prodrug peptidase sequences. We added our specific ZmaM sequence and aligned them with MAFFT^60,61^, removed sequences that were more than 80% identical, which resulted in a final alignment of 121 sequences. This alignment was composed of 55 type I and 66 type II prodrug peptidases. Logos were created using Weblogo 3^62^, and the sequence conservation was mapped onto the ZmaM structure using ConSurf^63^.

### List of accessorized ABC transporters

We extracted accessorized ABC transporter structures from the InterPro database^29^ accessed on July 15 2025. We searched by architecture domain, selecting the Pfam family PF00664 (ABC transporter transmembrane region) and curating the resulting domain architectures that include an additional Pfam family which comprises a transmembrane domain. These Pfam families included: PF24357 (TMD0_ABC), PF16185 (MTABC_N), PF24358 (ABCC10_N), PF00144 (β-lactamase). We curated this list to include only entries with experimental structures, which removed the β-lactamase architectures entirely. We further trimmed this list to include one structure of each conformation available for each individual protein, choosing the higher resolution structure if multiple were available. We manually included the structure of the transporter associated with antigen processing (PDB ID: 9RCV) as it also contains a membrane bound accessory domain but does not have an assigned Pfam entry. We aligned all the structures to ZmaM’s structure in PyMOL, and calculated BSA values using the PDBePISA webserver^64^.

## Data availability

Atomic coordinates for the reported structure in this work have been deposited to the Protein Data Bank under accession number 12SJ. The map of the cryo-EM reconstruction has been deposited to the Electron Microscopy Data Bank under accession number EMD-76731. Source data for Figs. 1, 3, 5 and Extended Data Figs. 4 are available on Dryad (TBD).

## Acknowledgements

We acknowledge the early contributions of Janny Liao in cloning the ZmaM mutants; Elizabeth J. May, Heather M. Frank, Shamayeeta Ray, and Samuel P. Berry for technical assistance; and Ute Hellmich and Gaudet lab members for discussions. This work was funded, in part, by National Institutes of Health (NIH) grants R01GM120996 (R.G.), R35GM161633 (R.G.), R01AI164647 (D.Z.R), R21AI171308 (D.Z.R.), and R01CA208834 (E.P.B.), the National Science Foundation Graduate Research Fellowship Program grant DGE1745303 (G.E.Z.) and an HHMI James H. Gilliam Fellowship (J.A.V.). E.P.B is a Howard HHMI Investigator. M.D. acknowledges support from the Canadian Institutes of Health Research Postdoctoral Fellowship under grant no. MFE-176575. We acknowledge use of resources from the Harvard Cryo-EM Center for Structural Biology.

---

**Extended Data Table 1.** Cryo-EM data collection, refinement and validation statistics.

| **Data collection and processing** |  |
|---|---|
| Nominal Magnification | 165,000 |
| Voltage (kV) | 300 |
| Electron exposure (e<sup>-</sup>/Å<sup>-2</sup>) | 62.5/65.1 |
| Defocus range (µm) | 0.8, 2.2 |
| Pixel size (Å) | 0.825 |
| Symmetry imposed | C2 |
| Initial particle images (no.) | 2,159,824 |
| Final particle images (no.) | 1,081,062 |
| Map Resolution (Å) | 2.75 |
| FSC threshold | 0.143 |
| Map resolution range (Å) | 2.6-7.1 |
| **Refinement** |  |
| Initial model used | *ab initio* |
| Model resolution (Å) | 2.70 |
| FSC threshold | 0.500 |
| Map-sharpening B-factor (Å<sup>2</sup>) | 104.52 |
| Model composition |  |
| Non-hydrogen atoms | 15,730 |
| Protein residues | 1,952 |
| B factors (Å<sup>2</sup>) |  |
| Protein | 39.95 |
| Ligand | 13.95 |
| R.M.S.D. deviations |  |
| Bond lengths (Å) | 0.005 |
| Bond angles (°) | 0.75 |
| **Validation** |  |
| MolProbity score | 1.00 |
| Clashscore | 2.25 |
| rotamer outliers (%) | 0.18 |
| Ramachandran plot |  |
| Favored (%) | 98.15 |
| Allowed (%) | 1.85 |
| Disallowed (%) | 0 |

**Extended Data Figure 1.**
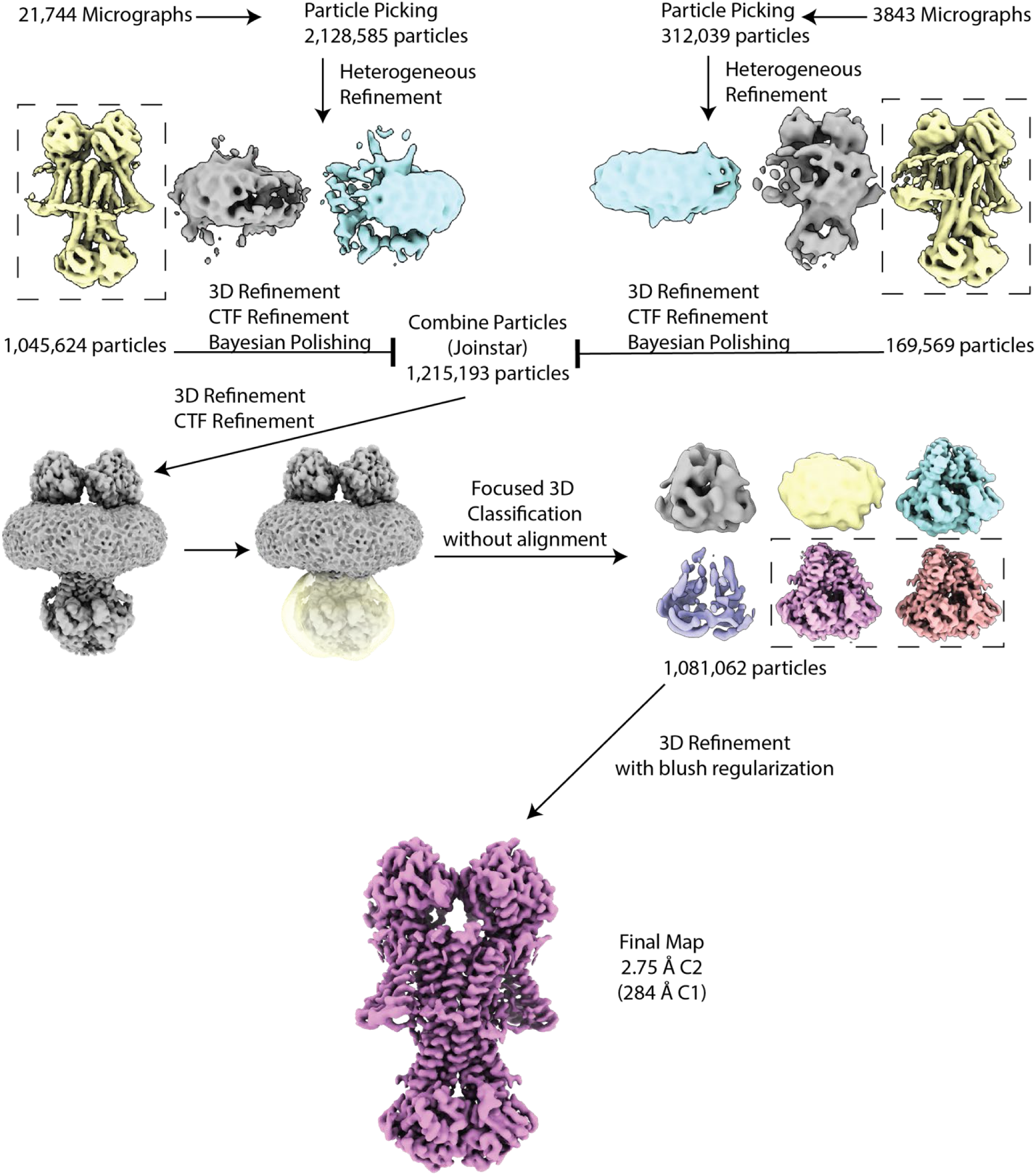
Cryo-EM processing workflow for the ZmaM E965Q structure. Details on the processing workflow for data collected for the E965Q structure. Locally filtered map with dust hidden used for the final reconstruction for clarity.

**Extended Data Figure 2.**
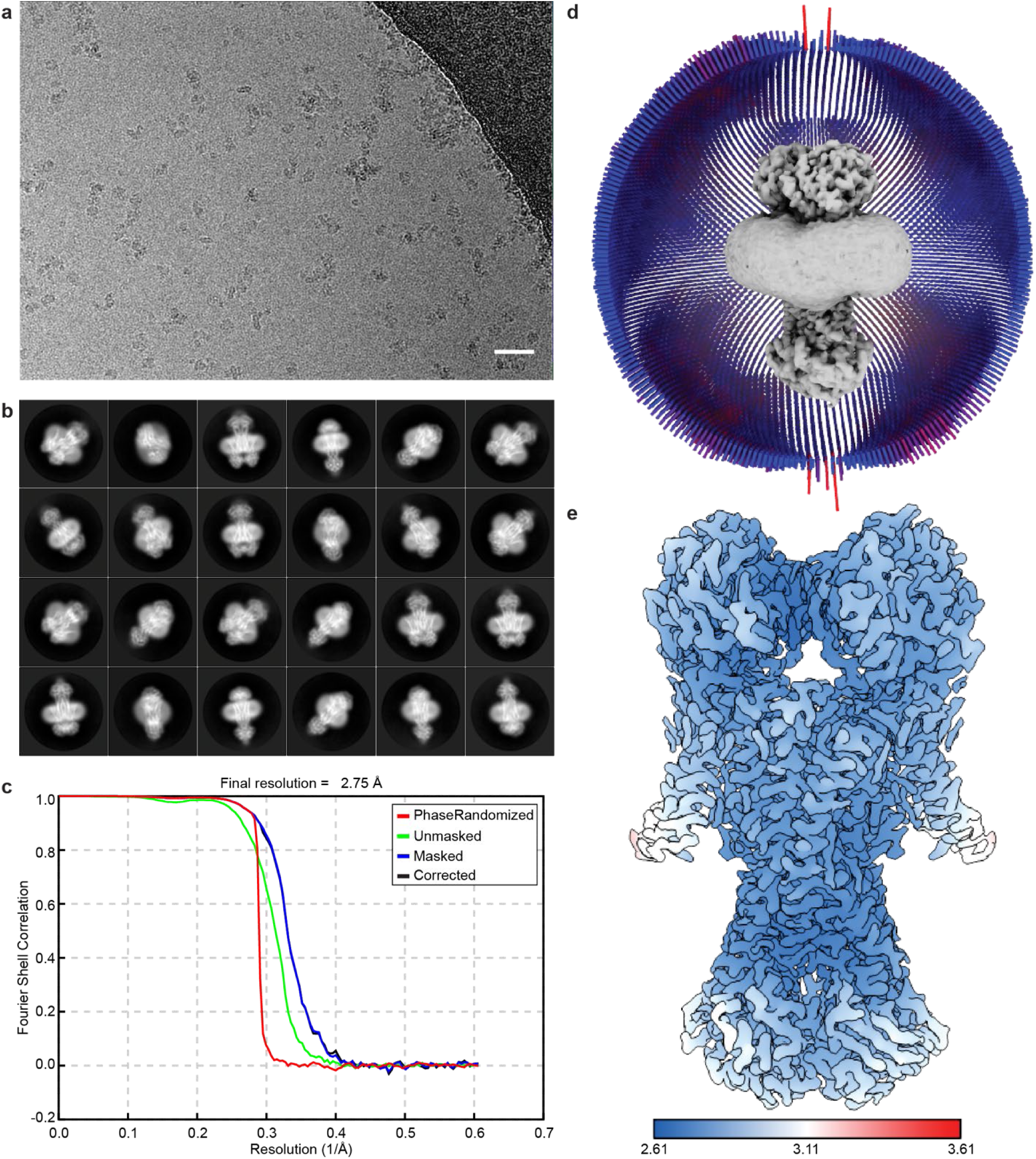
Cryo-EM map details for E965Q structure. a, Representative micrograph of 21,744 total micrographs from the ZmaM E965Q cryo-EM data collection, low-pass filtered for clarity. Scale bar = 250 Å. **b,** Selected 2D class averages. **c,** Gold-standard Fourier shell correlation (FSC) curves extracted from cryoSPARC. **d,** Angular distribution plot of all particles that contributed to the final map. **e,** Local resolution map. Color bar ranges from 2.61 Å to 3.61 Å.

**Extended Data Figure 3.**
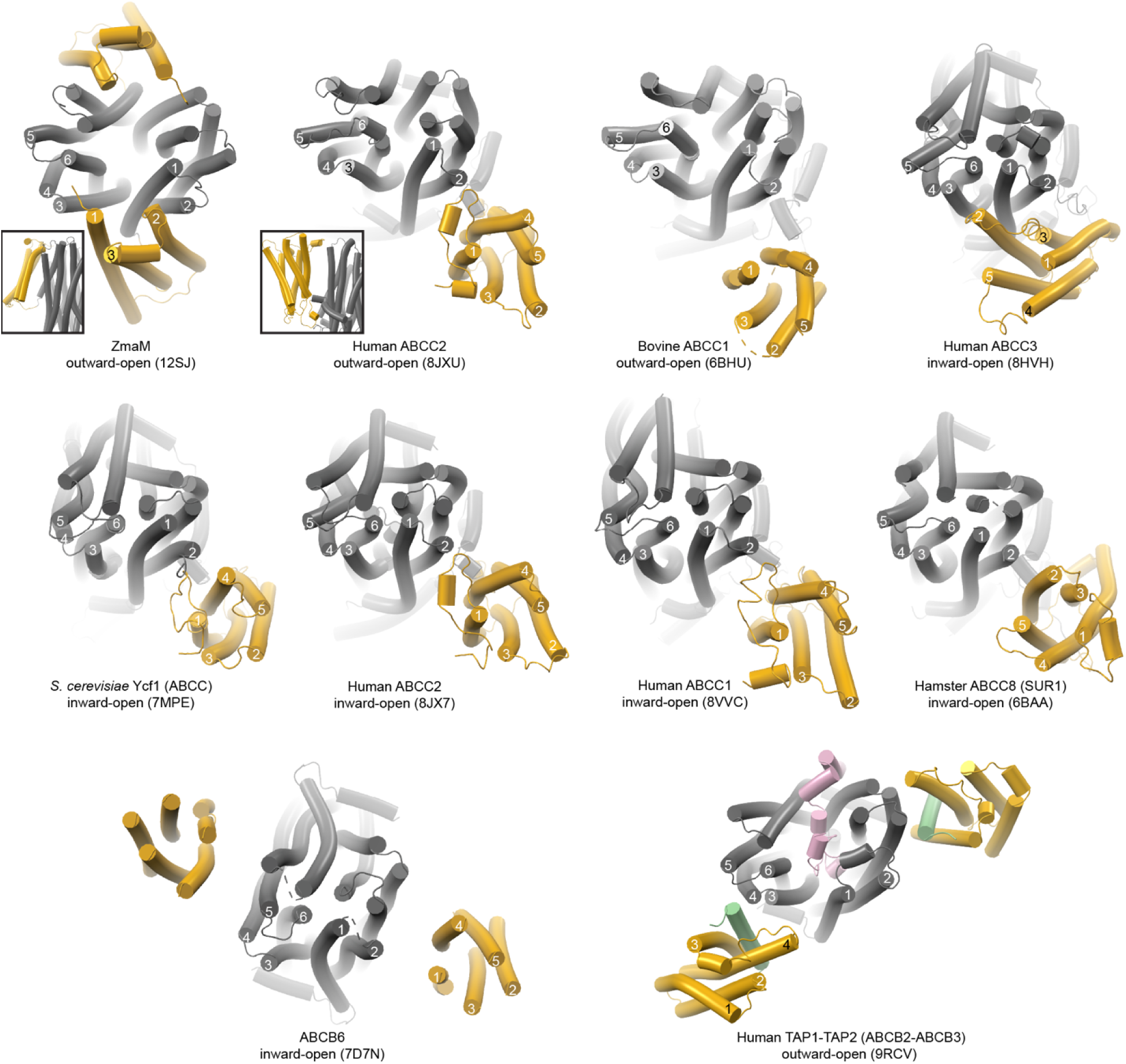
Superimposed structures of accessorized ABC transporters. All structures are superimposed relative to the ZmaM structure and viewed from the extracellular face at the level of the membrane outer leaflet. The core ABC transporter is shown as gray cylinders and the accessory domain as yellow cylinders. The human TAP1-TAP2 transporter structure (9RCV) includes US6 (pink) a TAP transporter inhibitory protein from human cytomegalovirus, and the transmembrane helix of tapasin (green). TMD numbering is included for one chain (or one half) of the transporter and accessory domain. The conformation captured in each structure and their respective PDB IDs are indicated below each structure.

**Extended Data Figure 4.**
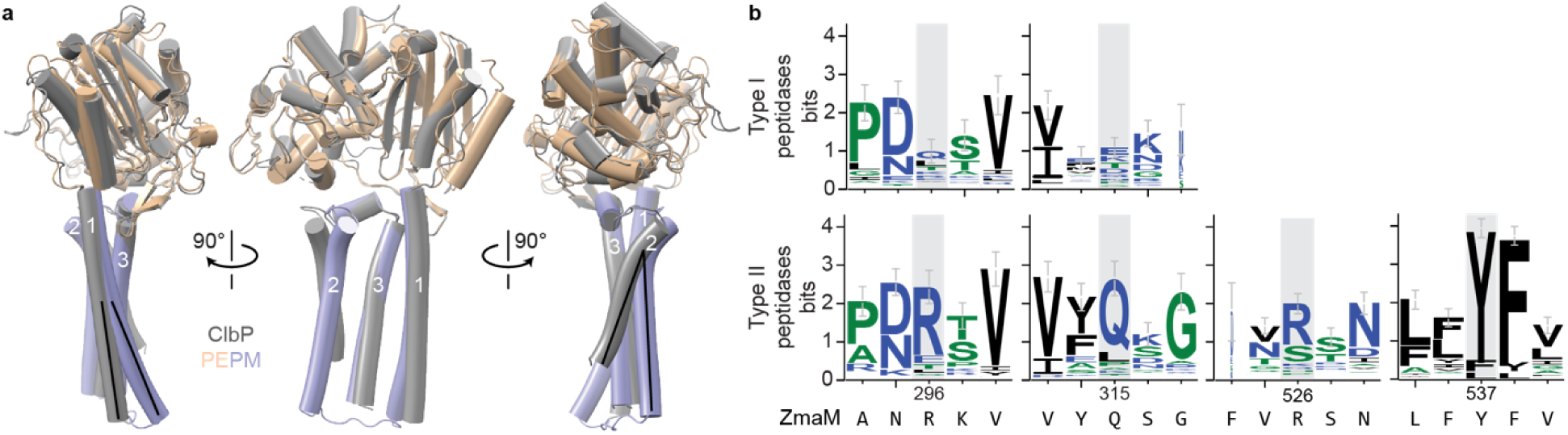
The ClbP and ZmaM peptidase modules have similar structures, and interface residues are conserved in type II prodrug peptidases. a, Structural alignment of ClbP (PDB ID: 7UL6, gray) and a peptidase module from ZmaM (wheat and light blue). TM helices are numbered and black lines highlight the pitch of distinct helices (see main text). b, Sequence logos representing conservation of, from left to right, ZmaM residues R296 and Q315 and their neighboring residues in alignments of 56 type I (top) or 67 type II (bottom) prodrug peptidases, and R526 and Y537 present only in type II peptidases. The corresponding ZmaM sequence is shown below the logo.

