## Supplementary Information for "Structure of *Bacillus cereus* ZmaM, a peptidase-ABC transporter that secretes and activates zwittermicin A"

This file contains:

- Supplementary Note 1. Cryptic transcription-translation in ZmaM
- Supplementary Tables 1-2
- Supplementary Figures 1-8

**Supplementary Note 1**. **Cryptic transcription-translation in ZmaM**

Purifications of ZmaM expressed in *E. coli* from its natural coding sequence included a consistent ~30-kDa contaminant detectable in anti-His-tag western blots, suggesting it contained a His-tag (Supplementary Figs. 5 and 8a). This contaminant also appears in un uninduced samples but not empty-vector samples (Supplementary Fig. 8a), further suggesting that it arises from a promoter other than the inducible T7lac promoter preceding the *zmaM* gene. Chymotrypsin digestion and mass spectrometry of the isolated gel band indicated that it corresponds to a C-terminal fragment of ZmaM starting after V742 at N743 (Supplementary Fig. 8b-c). However, chymotrypsin cleaves C-terminal to Tyr, Phe, Trp, Leu, and Met residues, but not after Val residues. The V742 codon is GTG, which can serve as a non-canonical start codon for *E. coli* translation. Translation initiating at this site, followed by removal of the N-terminal methionine would explain the observed contaminant. There is also a nearby ATG codon (M744) and a sequence resembling the consensus *E. coli* Shine-Dalgarno sequence upstream from the V742 codon (consensus: AGGAGGU, *zmaM*: CGGAGAU).

To test this hypothesis and address the contaminant's presence, we introduced a series of mutations. We mutated the alternate start V742 codon GTG to GTT (synonymous mutation) and the canonical start M744 codon ATG to CTG (mutating Met to Leu). Of note, M744 is exposed to solvent in the extracellular loop between TMs 5 and 6 and its sidechain density is not resolved in the structure, suggesting that the impact of an M744L mutation is minimal. Position 744 is generally conserved as an aliphatic residue in type II prodrug peptidases (Supplementary Fig. 8d). We also introduced additional silent mutations in the hypothetical Shine-Dalgarno sequence (to CAGAAAT) to distinguish it further from the consensus sequence. Each mutation reduced the amount of truncated product (Supplementary Fig. 8e). We combined all these mutations to minimize the amount of ~30-kDa product produced and moved the His-tag to the ZmaM N-terminus to minimize copurification of the ~30-kDa product with His-tagged ZmaM by Ni-NTA affinity chromatography. Comparing the purity of N-terminally His-tagged ZmaM from the reengineered sequence to that of C-terminally His-tagged ZmaM produced from its natural coding sequence, the amount of contaminant was greatly reduced (Supplementary Fig. 8f). To minimize the potential effects of this truncated ZmaM on *in vitro* activity measurements, we used this modified ZmaM as the "WT*" in all *in vitro* functional analysis experiments.

**Supplementary Table 1. Curated list of accessorized ABC transporters.**

| PDB ID | Name | Resolution (Å) | Proteins | Conformation | BSA ($Å^2$) |
|---|---|---|---|---|---|
| 12sj | ZmaM | 2.8 | NA | Outward-open | 564 |
| N/A | ZmaM AF model | N/A | NA | Occluded | 849 |
| N/A | ZmaM AF model | N/A | NA | Inward-open | 980 |
| 6bhu | Bovine ABCC1 | 3.14 | q8hxq5 | Outward-open | n.d.[a] |
| 8vvc | Human ABCC1 | 4.32 | p33527 | Inward-open | 0 |
| 8jxu | Human ABCC2 | 3.55 | q92887 | Outward-open | 26 |
| 8jx7 | Human ABCC2 | 3.6 | q92887 | Inward-open | 20 |
| 8hvh | Human ABCC3 | 3.07 | o15438 | Inward-open | 885 |
| 6baa | Hamster ABCC8 | 3.63 | q09427 | Inward-open | 178 |
| 7mpe | *S. cerevisiae* Ycf1 | 3.2 | p39109 | Inward-open | 172 |
| 9rcv | Human ABCB2-ABCB3 | 2.7 | q03518 | Outward-open | 44 |
| 7d7n | Human ABCB6 | 5.2 | q9np58 | Inward-open | 0 |

[a]Not determined. Only the backbone of the accessory domain is modeled, precluding measurement of the interface BSA.

**Supplementary Table 2. Cloning primers.**

| Primer Name | Sequence 5' – 3' |
|---|---|
| **Z UP F** | AGCGTACTCGGTCGACGACTCGGTATTGGTTTCGATTG |
| **Z UP R** | GCCTCCTTTGTTTTTAACTTCATATTTTTAGCCCTCTTTTAG |
| **Z DOWN F** | GAAGTTAAAAACAAAGGAGGCATTATTATGAGTAAG |
| **Z DOWN R** | GGACGAATTATACAAATAAGGATCCCTAAAATTAATATTCCATTTTCGTTGCTAGTTTC |
| **C UP F** | GCTTGCATGCCTGCAGGTCGACGAGATGAAGCAAGAGGTGC |
| **C UP R** | GTGTATGAAAAAGTTAGATGCCTCGTTCCG |
| **C DOWN F** | GCATCTAACTTTTTCATACACTCTCCTTTCTGAAG |
| **C DOWN R** | GGACGAATTATACAAATAAGGATCCGACCCCAACGAAAAACAAAAC |
| **PulA UP F** | GAGCGCAGCGAGTCAGTGAGCGAGGAAGCGGAAGAGCGTAATAACTCTTGTGCGTGAAGC |
| **PulA UP R** | GTTTTCTTTAAGACGATAATGTGACAAATCGCTACTTG |
| **PulA DOWN F** | CATCATTTCCTTCCTTTTACTTTCAACATAAAAAAATCACCATCATTCTC |
| **PulA DOWN R** | CGACGTTGTAAAACGACGGCCAGTGAATTCCATATTACCCTTTCCTTTTTAGGAGG |
| **ZmaM_ins_F** | GCTTACATAAGGAGGAACTACTATGAAGTTAAACATATGGTTGAAGTTTATC |
| **ZmaM_ins_R** | GCATGCGAGGTGCGGCCTCAGTCAATGATGGTGATGGTGGTG |
| **BGC_Pro_F** | GTCACATTATCGTCTTAAAGAAAACTTCTTGTGAATATGTTACTAACTAC |
| **BGC_Pro_R** | CTTATGTAAGCTTATTTTGTTTAATTCAGATTTTCTATGCTTTAC |
| **ZmaM F** | GAAGGAGATATACATATGAAGTTAAACATATGGTTGAAGTTTATCATTTTA |
| **ZmaM R** | GTGATGGTGATGATGAGCGGCCGCTAATAATGCCTCCTTTGTTTTATTCATTTTCAC |
| **S89A_F** | GCTAGGTGCTACTTCAAAAGCTTTTACGGCATTAGCTG |
| **S89A_R** | GAAGTAGCACCTAGCTCGAAAAGCGTCTCATTCG |
| **R296A_F** | CGAATGCGAAGGTTGCCCCAGATAACAATGGAGCTTCATATGCAG |
| **R296A_R** | CAACCTTCGCATTCGCTTCATGAGAATCGTTAATTAATTCCTCCGGAAACATC |
| **R296E_F** | CGAATGAAAAGGTTGCCCCAGATAACAATGGAGCTTCATATGCAG |
| **R296E_R** | CAACCTTCTTATTCGCTTCATGAGAATCGTTAATTAATTCCTCCGGAAACATC |
| **Q315A_F** | GTCTATGCGAGTGGTGGAGGGGAAATTTCACATGGTGGAAGTAACC |
| **Q315A_R** | CCACTCGCATAGACCTCCCATCCAGCTGCATATGAAGCTCC |
| **R526A_F** | CGTTTGTTGCATCAAATAATTTGGAAAATGGCTTGTTGTTTTACTTTGTACTTGGAATCG |
| **R526A_R** | CCAAATTATTTGATGCAACAAACGTTTCATTAATCACAAAAATAAGAAAGGCATTACC |
| **R526E_F** | CGTTTGTCGAAAGCAATAATCTGGAGAATGGTCTGCTGTTCTATTTCGTGC |
| **R526E_R** | ATTGCTTTCGACAAACGTTTCGTTGATCACGAAGATCAAGAAGGCATTACCAAAG |
| **Y537A_F** | CTGTTCGCGTTCGTGCTGGGTATTGTTATGTATGTGTTTAGCCAACG |
| **Y537A_R** | CACGAACGCGAACAGCAGACCATTCTCCAGATTATTGCTACGG |
| **E965Q_F** | TTGATCAATGGGCAGCGGATCAAGATCCTGGTTATAG |
| **E965Q_R** | CGCTGCCCATTGATCAAATAAAAGTATAGGTCGATCTTCC |
| **ABCM_F** | GATATACTTATGCATCATCACCATCACCATCACCACAATCTTTGTTTTCCAAGACAAAAG |
| **ABCM_R** | GTGGTGGTGGTGCTCGAGTGCCACTATTATAATAATGCCTCCTTTG |
| **PEPM_F** | GAAGGAGATATACATATGAAGTTAAACATATGGTTGAAGTTTATCATTTTA |
| **PEPM_R** | GTGATGGTGATGATGAGCGGCCGCCGACTTTCCCTTTTGTCTTGG |
| **SD Mut F** | CCCAGAAATCTTTGTGAATATGCCGAAAGAAACCATCAGC |
| **SD Mut R** | CAAAGATTTCTGGGAAAACAAACACCACAATACCGATGAC |
| **Start Cod F** | CTTTGTTAATCTGCCGAAAGAAACCATCAGCAACTATGTGATCGTTTTC |
| **Start Cod R** | CGGCAGATTAACAAAGATCTCCGGGAAAACAAACACCAC |
| **SD Start F** | CCCAGAAATCTTTGTTAATCTGCCGAAAGAAACCATCAGC |
| **SD Start R** | CGGCAGATTAACAAAGATTTCTGGGAAAACAAACACCAC |

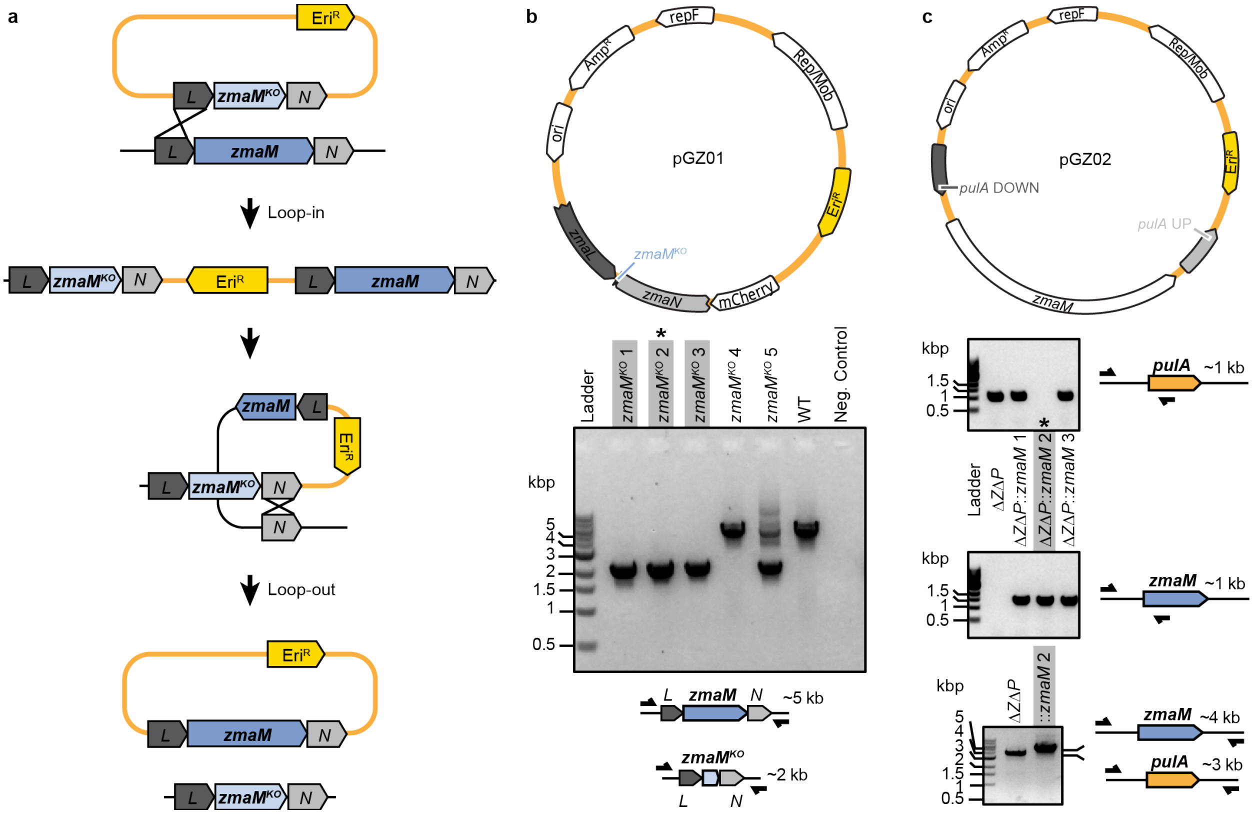


**Supplementary Figure 1. Approach to *B. cereus* genetic engineering. a,** General workflow of *B. cereus* loop-in-loop-out cloning with the plasmid (yellow track) harboring the erythromycin resistance cassette ($Eri^R$, yellow), the truncated ZmaM coding sequence ($zmaM^{KO}$, light blue), and the upstream and downstream homology regions *zmaL* (*L*, dark gray) and *zmaN* (*N*, light gray). The black track represents the *B. cereus* genome with the full-length *zmaM* gene (blue). **b,** Map of plasmid used to introduce $zmaM^{KO}$ into *B. cereus* (to) and the agarose gel confirming successful loop-in-loop-out in 3 of the 5 tested clones by PCR amplification from the strains' genomic DNA. Schematics on the bottom illustrate amplicon size difference based on the gene's identity. Gray boxes mark the strains that successfully incorporated the truncated gene. Clone 2 (marked with an asterisk) was then used as the Δ*zmaM* strain throughout this study. **c,** Map of plasmid used to reintroduce *zmaM* gene into the *B. cereus pulA* locus and the agarose gels confirming successful insertion by PCR amplification from the strains' genomic DNA. Top and middle gels show products from PCR reactions that selectively amplify based on the identity of the gene present at the locus at the reverse primer's binding site, as shown in the schematic to the right. Bottom gel shows products from PCR reactions that amplify the entire insert region, with the product size dependent on the gene present at the locus. Gray boxes and an asterisk mark the strain that successfully incorporated the *zmaM* gene.

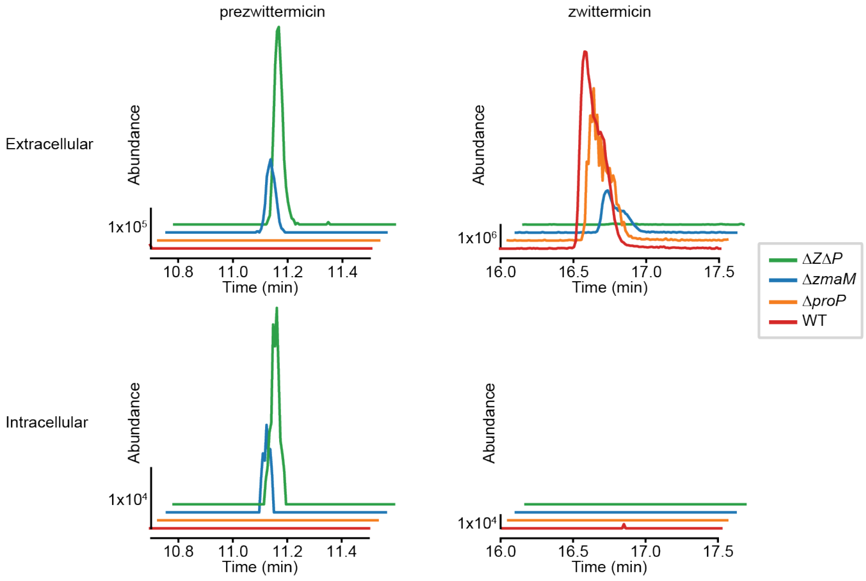


**Supplementary Figure 2. LC-MS analysis of Δ*proP*, Δ*zmaM*, and Δ*Z*Δ*P*.** LC-MS traces of extracellular (top) and intracellular (bottom) prezwittermicin (left) and zwittermicin (right) from WT, Δ*proP*, Δ*zmaM*, and Δ*Z*Δ*P* strains. These are representative traces from one of two independent replicate experiments.

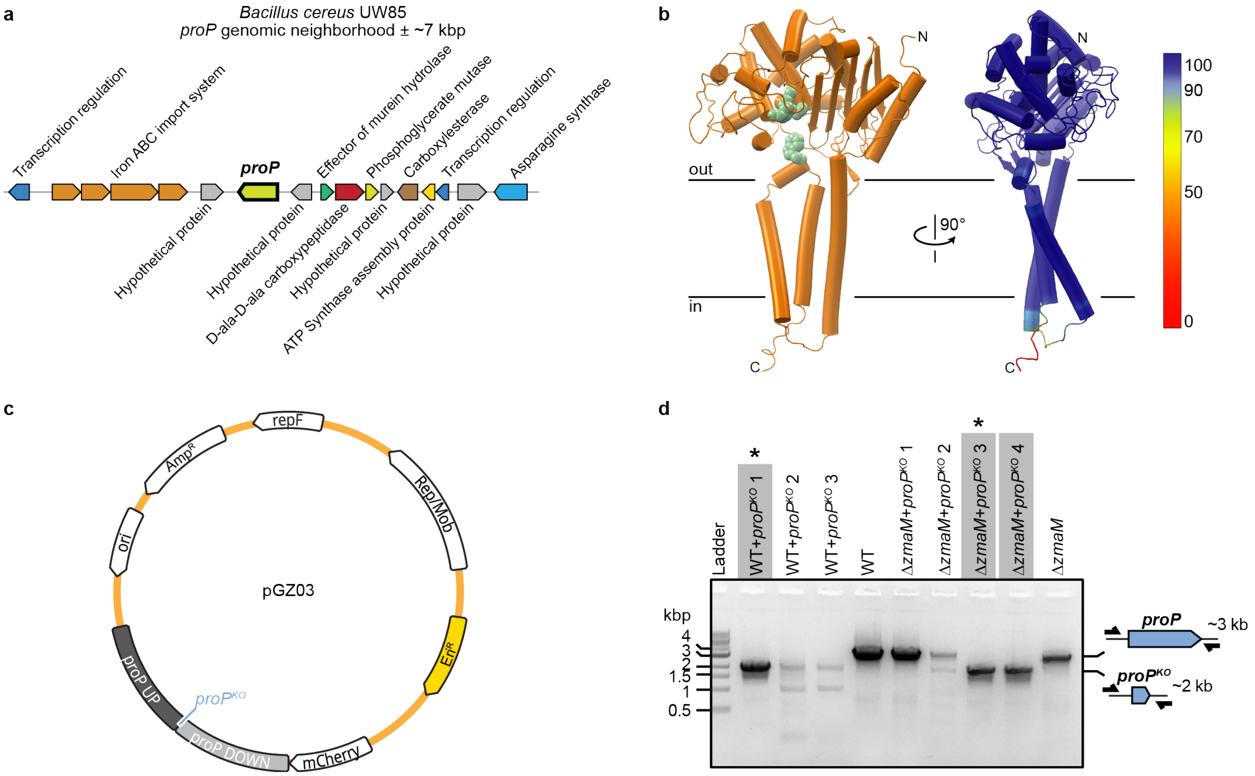


**Supplementary Figure 3. Genomic neighborhood, predicted structure, and gene knockout of ProP. a,** Genomic neighborhood of *proP* (± ~7 kbp) in the *B. cereus* UW85 genome with genes annotated by the JGI IMG database[1]. **b,** Cartoon representations of the Alphafold model of ProP. On the left, the conserved catalytic triad and tryptophan are highlighted as green spheres. The view to the right is rotated by 90° and colored by per-residue pLDDT confidence score. **c,** Map of the plasmid used for loop-in/loop-out to knock-out the *proP* gene in *B. cereus*. **d,** Agarose gel of PCR amplicons confirming successful truncation of the *proP* gene by loop-in/loop-out in 1 of 3 WT+*propP*$^{KO}$ and 2 of 4 Δ*zmaM*+*propP*$^{KO}$ clones tested, respectively. The schematics on the right highlight amplicon sizes of the wildtype *proP* and truncated *proP*$^{KO}$ genes. The successful clones are boxed in grey; WT+*propP*$^{KO}$ clone 1 and Δ*zmaM*+*propP*$^{KO}$ clone 3 (both marked with asterisks) were used as strains Δ*proP* and ΔZΔ*P*, respectively, throughout this study.

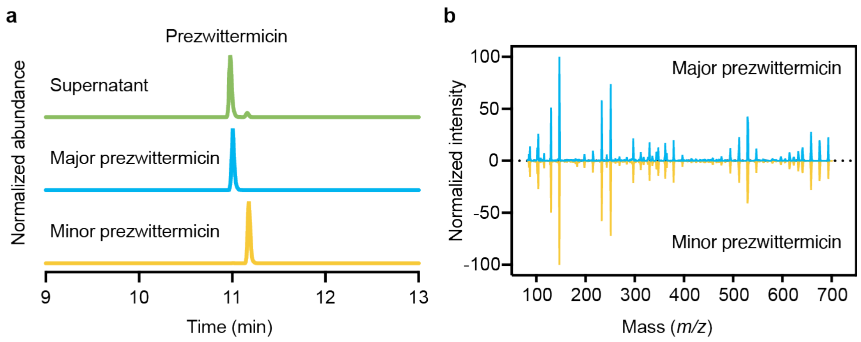


**Supplementary Figure 4. Prezwittermicin is produced as a mixture of isomers. a,** EICs ($[M+H]^+$ = 693.4141 *m/z*) of the spent supernatant from *B. cereus* and from the purified major and minor prezwittermicins. The intensity is normalized to the maximum value in each sample. **b,** MS/MS fragmentation results of the purified major and minor prezwittermicins. The intensity is normalized to the maximum value in each sample.

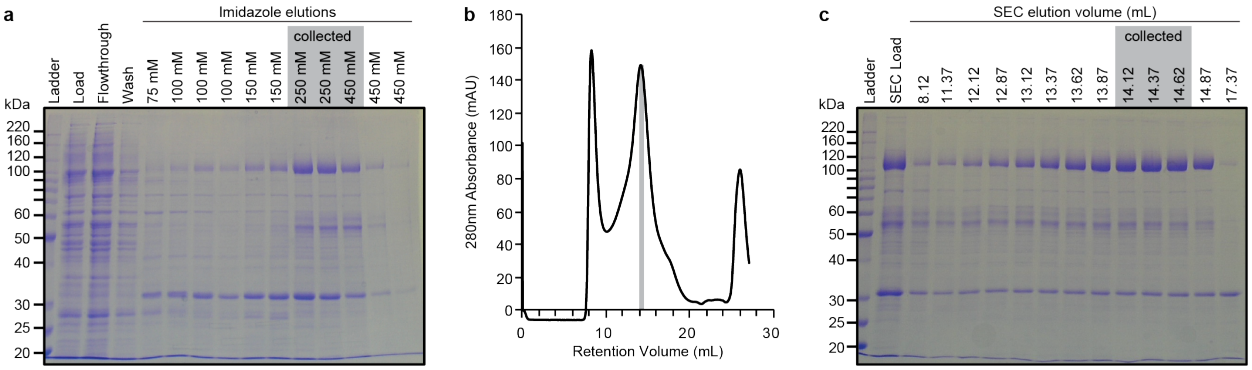


**Supplementary Figure 5. ZmaM E965Q purification. a,** Coomassie-stained SDS-PAGE gel evaluating the nickel-immobilized metal affinity chromatography of ZmaM E965Q, showing the load, flowthrough, wash, and imidazole-based elution fractions. Fractions collected for further purification are boxed in grey. **b,** Size-exclusion chromatogram for ZmaM E965Q purification. The gray box indicates fractions collected for cryo-EM grid preparation. **c,** Coomassie-stained SDS-PAGE gel evaluating the SEC load and elution fractions from the chromatogram in **b**. The gray box indicates the fractions used for cryo-EM grid preparation.

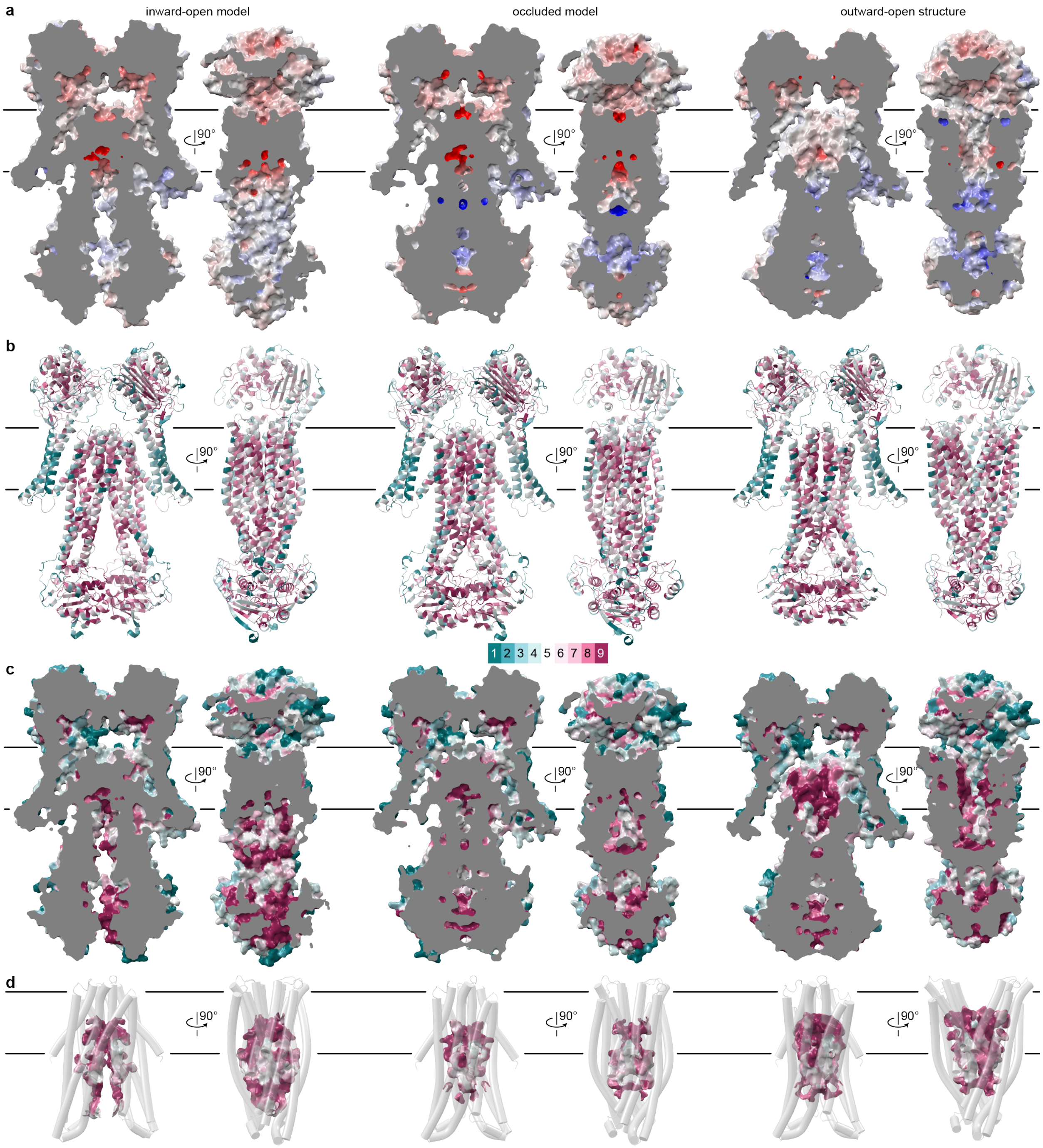


**Supplementary Figure 6. Sequence conservation in type II prodrug peptidases.** All panels show, from left to right, the inward-open model, occluded model, and outward-open structure of ZmaM, including head-on (left) and side views (right; front peptidase module omitted to better visualize the transporter module). **a,** Cross-section of the surface electrostatics as predicted by APBS[65], from -20 (red) to +20 kcal/mol (blue). **b,** Cartoon representation with esidues colored by ConSurf score from cyan (variable) to magenta (conserved). **c,** Cross-section of the surface representation colored by ConSurf scores. **d,** Transporter module TMD as transparent cartoon with surface representation of the cavities colored by ConSurf score.

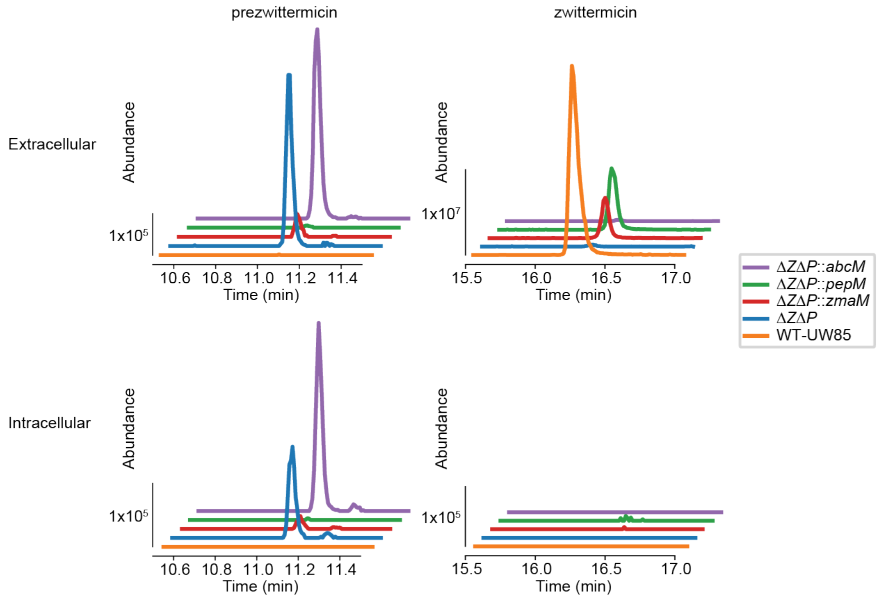


**Supplementary Figure 7. LC-MS analysis of Δ*Z*Δ*P* *trans*-complemented with ZmaM constructs.** LC-MS traces of intracellular and extracellular prezwittermicin (EIC for $[M+H]^+$ = 693.4141) and zwittermicin (EIC for $[M+H]^+$ = 315.2278) from WT, Δ*Z*Δ*P*, and Δ*Z*Δ*P* *trans*-complemented with ZmaM and the half constructs PepM and ABCM. These are representative traces from one of two independent replicate experiments.

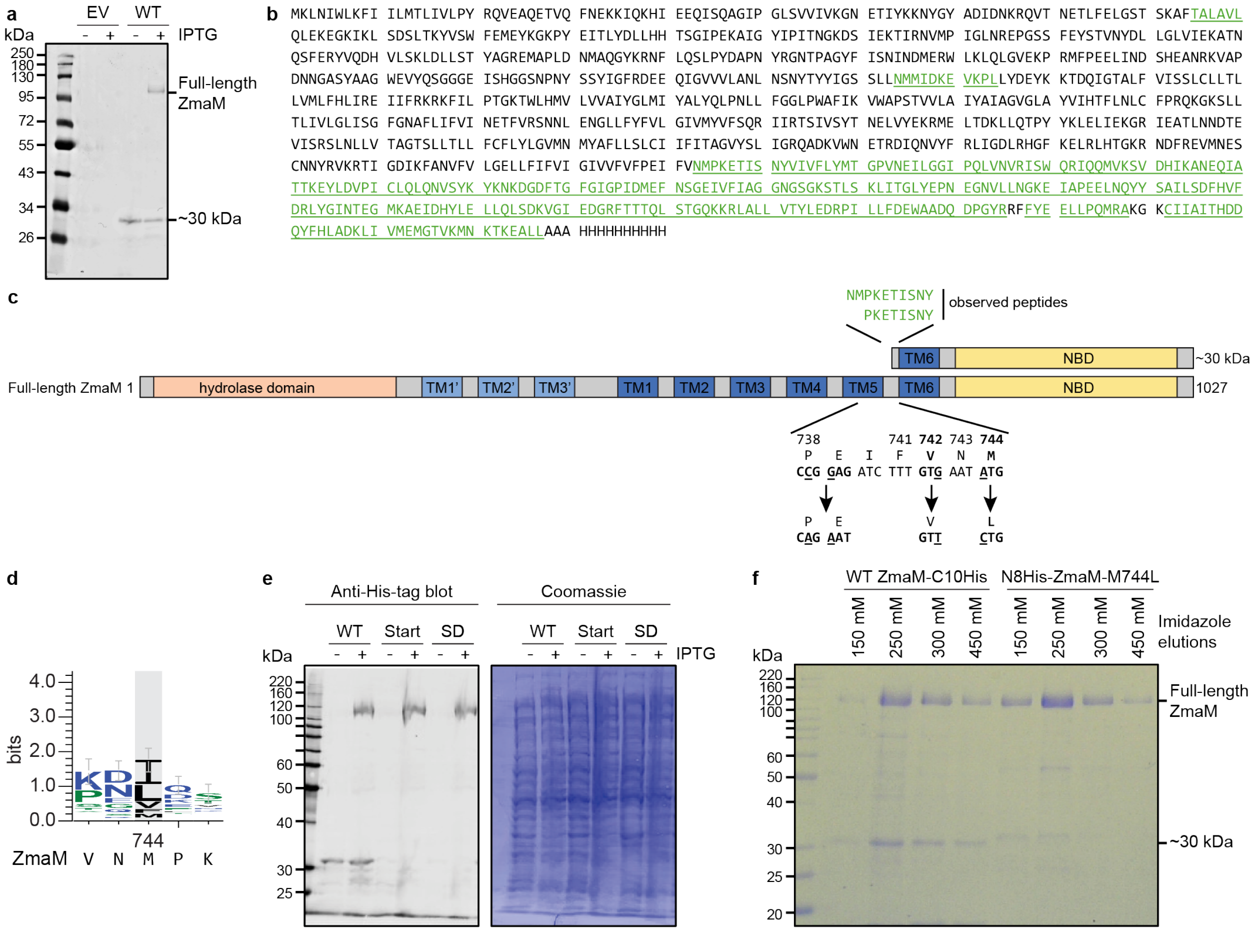


**Supplementary Figure 8. Identification and mitigation of a ~30-kDa contaminant arising from cryptic transcription and translation of the WT ZmaM open reading frame expressed in *E. coli*. a,** Anti-His western blot of uninduced or isopropyl β-D-1-thiogalactopyranoside (IPTG)-induced whole-cell lysates of *E. coli* transformed with empty vector (EV) or expressing wildtype (WT) ZmaM. **b,** ZmaM sequence highlighting in green the N-terminal sequencing mass spectrometry peptide reads from the gel-purified ~30 kDa contaminant in purifications of C-terminally His-tagged wildtype ZmaM. Essentially all peptides map to C-terminus of ZmaM, with the most N-terminal peptides from that ~30 kDa C-terminal fragment starting at N743 or P745 (see also panel c). A fragment starting at N743 corresponds to a theoretical molecular mass of 34.1 kDa. **c,** Schematic of the full-length ZmaM domain structure and identified ~30-kDa contaminant. The two most N-terminal peptides from the contaminant observed by mass spectrometry are shown above. Below is relevant region of the ZmaM protein and coding sequences, with the mutations introduced to reduce the amount of co-purified ~30-kDa contaminant. **d,** Sequence logo of positions 742-746 (ZmaM numbering) across 67 aligned type II prodrug peptidase sequences. The corresponding ZmaM sequence is shown below the logo. **e,** Anti-His western blot (left) and corresponding Coomassie-stained blot (right) of uninduced or IPTG-induced whole-cell lysates of *E. coli* expressing either wildtype (WT) ZmaM, or variants with mutations of the start codons (Start) or potential internal Shine-Dalgarno (SD) sequence. **f,** Coomassie-stained SDS-PAGE gel comparing the purity of nickel-immobilized affinity chromatography elutions for WT ZmaM with a C-terminal 10xHis tag (left) and ZmaM with an N-terminal 8x-His tag and the mutations illustrated in panel c (right).
